# Prediction of subgenome additive and interaction effects in allohexaploid wheat

**DOI:** 10.1101/373605

**Authors:** Nicholas Santantonio, Jean-Luc Jannink, Mark E. Sorrells

## Abstract

Whole genome duplications have played an important role in the evolution of angiosperms. These events often occur through hybridization between closely related species, resulting in an allopolyploid with multiple subgenomes. With the availability of affordable genotyping and a reference genome to locate markers, breeders of allopolyploids now have the opportunity to manipulate subgenomes independently. This also presents a unique opportunity to investigate epistatic interactions between homeologous orthologs across subgenomes. We present a statistical framework for partitioning genetic variance to the subgenomes of an allopolyploid, predicting breeding values for each subgenome, and determining the importance of inter-genomic epistasis. We demonstrate using an allohexaploid wheat breeding population evaluated in Ithaca, NY and an important wheat dataset previously shown to demonstrate non-additive genetic variance. Subgenome covariance matrices were constructed and used to calculate subgenome interaction covariance matrices across subgenomes for variance component estimation and genomic prediction. We propose a method to extract population structure from all subgenomes at once before covariances are calculated to reduce collinearity between subgenome estimates. Variance parameter estimation was shown to be reliable for additive subgenome effects, but was less reliable for subgenome interaction components. Predictive ability was equivalent to current genomic prediction methods. Including only inter-genomic interactions resulted in the same increase in accuracy as modeling all pairwise marker interactions. Thus, we provide a new tool for breeders of allopolyploid crops to characterize the genetic architecture of existing populations, determine breeding goals, and develop new strategies for selection of additive effects and fixation of inter-genomic epistasis.

## 2 Introduction

Gene duplication is known to be a primary driver of evolution by providing the raw genetic material for gene diversification through sub- and neofunctionalization (Haldane 1933; Ohno 1970). Whole genome duplication events, in which an entire set of genes is duplicated, occurs either through duplication of the same genome (autopolyploidy) or the union of two closely related genomes (allopolyploidy). Both types of polyploids can exhibit non-additive genetic variation from the presence of multiple alleles (Segovia-Lerma et al. 2004; Birchler et al. 2010; Jiang et al. 2017), although how these non-additive effects are classified needs clarification.

Statistical deviations from additivity (i.e. interactions) are important contributors to genetic variation. Homologous gene interactions, also known as dominance, are deviations from an additive expectation due to different allele combinations at one locus. Non-homologous gene interactions, commonly referred to as epistasis, are deviations from an additive expectation due to different allele combinations at two or more loci (Fisher 1919). When epistasis occurs between non-homologous loci with similar function, such as across orthologs or paralogs, these interactions are comparable to dominance effects. If interactions occur between homeologous orthologs on separate subgenomes of an allopolyploid, should we call this epistasis or dominance?

In classical hybrid variety production, divergent sets of alleles are intentionally isolated into heterotic groups and then brought back together to form a hybrid. This establishes heterozygosity (by decent) at all loci to form a homogeneous population. The union of two divergent suites of genes during the formation of an allopolyploid also results in a homogeneous population, but heterozygosity is established across homeologs rather than homologs. Diploid hybrids lose heterozygosity through segregation in following filial generations, but heterozygosity across homeologous genes is subsequently preserved through selfing in the allopolyploid (Mac Key 1970; Ozkan, Levy, and Feldman 2001; Abel, Möllers, and Becker 2005). Allelic interactions contribute to dominance variance in the diploid hybrid, whereas homeoallelic interactions will be present as part of the additive by additive epistatic variance in an inbred allopolyploid population. As such, allopolyploids may be thought of as an immortalized hybrid (Ellstrand and Schierenbeck 2000; Feldman et al. 2012), although it is not yet clear that these exhibit a true heterotic response as traditional hybrids have demonstrated.

Birchler et al. (2010) note that newly synthesized allopolyploids often outperform their sub-genome progenitors, and that the heterotic response appears to be exaggerated in wider inter-specific crosses. This seems to hold true even within species, where autopolyploids tend to exhibit higher vigor from wider crosses (Bingham et al. 1994; Segovia-Lerma et al. 2004). Complementation of deleterious recessive alleles (or pseudo-dominance) has long been the primary explanation of the heterotic response (Stuber et al. 1992; Cockerham and Zeng 1996). However, Birchler et al. (2010) indicate evidence against this, where purging deleterious alleles has increased the additive value of inbred parents but has not reduced the heterotic response observed in the hybrid (Duvick 1999). Complementation also seems an unlikely driver of a heterotic response in allopolyploids, as the inbred subgenome progenitors would supposedly need functional copies of these genes to survive.

The availability of affordable genome-wide markers has sparked a revolution in selection on additive variation through the use of genomic prediction models. The additive genetic merit of an individual can be estimated as the sum of its additive marker effects to produce a genomic estimated breeding value (GEBV) (Meuwissen, Hayes, and Goddard 2001). When the number of markers is large, marker effects are typically considered random and normally distributed such that only one parameter need be estimated. Alternatively, the additive genetic covariance between individuals can be estimated from the same genome-wide markers and used to predict additive genetic values of individuals based on relatedness (Nejati-Javaremi, Smith, and Gibson 1997; VanRaden 2008). These models are equivalent for prediction under the same set of assumptions (Garrick 2007; VanRaden 2008; Strandén and Garrick 2009). Genomic prediction models have since become popular for their ability to predict the performance of genotyped individuals with no phenotypic observations. Selections on unobserved individuals allows for reduction in the cost of phenotyping and breeding cycle time, increasing the rate of genetic gain (Goddard and Hayes 2007; Heffner, Sorrells, and Jannink 2009; Jannink, Lorenz, and Iwata 2010; Heslot, Jannink, and Sorrells 2015).

The potential utility of genome-wide markers has also drawn renewed interest in non-additive genetic variation in recent years (Vitezica, Varona, and Legarra 2013; Martini et al. 2016; Jiang and Reif 2015; Huang and Mackay 2016; Jiang et al. 2017). Genomic prediction models that use genome-wide markers can incorporate non-additive genetic components to obtain better estimates of individual performance than based on additivity alone (Technow et al. 2012; Vitezica, Varona, and Legarra 2013; Jiang and Reif 2015; Akdemir and Jannink 2015; Akdemir, Jannink, and Isidro-Sánchez 2017; Wolfe et al. 2016). In outcrossing species such as maize, prediction of dominance effects is key to harnessing heterosis in unobserved hybrids (Technow et al. 2012). In inbred species, additive byf additive epistatic effects have been shown to significantly increase genomic prediction accuracy (Crossa et al. 2010; Martini et al. 2016). Epistatic effects can be added to the prediction model by extending Henderson’s (1985) method of expected epistatic covariance estimation to marker based covariance estimation (Jiang and Reif 2015; Martini et al. 2016).

The use of genome-wide markers has allowed for the partitioning of genetic variance to specific units of chromatin, previously infeasible with phenotypes alone (Bernardo and Thompson 2016). Allopolyploids have been traditionally treated as diploids because they undergo disomic inheritance (Mac Key 1970), such that the contribution of each subgenome to the genetic variance is ignored. By assigning markers to each subgenome, an additive genetic covariance based on each subgenome can be calculated. Using these covariances in a genomic prediction model, the genetic merit of an allopolyploid individual can be assigned to each of its subgenomes. These subgenomic estimated breeding values (SGEBV) can then be used to identify parents with complementary subgenome effects for crossing.

Under Hardy Weinburg equilibrium, subgenomes segregate independently, and realized estimates of additive covariance of individuals based on each subgenome will be independent. However, this does not generally hold true in breeding programs, where population structure from non-random mating is inherent. As a consequence, the estimates of additive covariance between individuals based on different subgenomes will not be independent, potentially leading to confounding of effects from each subgenome and problems partitioning variance reliably. In an attempt to circumvent this obstacle, we present an approach for removing the largest sources of genetic variance (i.e. population structure) using singular value decomposition of the matrix of marker scores.

Common wheat (*Triticum aestivum*) is a staple allopolyploid crop which has undergone two allopolyploid events, resulting in three genomes, denoted A, B and D. The A genome ancestor, *Triticum uratu*, still exists today and was an early domesticate from the fertile crescent important in the neolithic revolution (Dvořák et al. 1993). The B genome ancestor (an *Aegilops spp*.) is believed to have since gone extinct (Blake et al. 1999), but the tetraploid formed by these two genomes, *Triticum turgidum*, is still cultivated today primarily as emmer wheat. The D genome comes from a goat grass, *Aegilops tauschii*, which was may have been incorporated in a single hybridization event as recently as 10,000 years ago (Salamini et al. 2002). However, recent evidence based on sequence divergence of the D genome from the A and B genome has suggested a much earlier D genome incorporation around 400,000 years ago (Marcussen et al. 2014). Other evidence shows that limited gene flow into the D genome may have occurred after the polyploidization event, but appears to be from a single lineage (Wang et al. 2013). As a result, the D genome has significantly lower genetic variation than either the A or B genome.

We demonstrate this methodology using two allohexaploid wheat data sets, the Cornell small grains breeding program soft winter wheat breeding population dataset presented here (CNLM) and the W-GY wheat data from Crossa et al. (Crossa et al. 2010).

## 3 Materials and Methods

### 3.1 Empirical data sets

#### 3.1.1 CNLM population

The CNLM dataset consists of 8,692 phenotypic records of 1,447 soft winter wheat inbred lines evaluated at four locations near Ithaca, NY from 2007 to 2016, representing 26 environments (Table A1). These phenotypic evaluations serve primarily as a first round of selection for grain yield and other agronomic traits before relatively few are selected for replicated regional trials around New York State. Lines are introduced and then removed after they are deemed either fit for advanced field trials or to be discarded or recycled in the breeding program. As such, this dataset is unbalanced in nature. Most lines were not replicated within a given trial (i.e. year and location), but various check varieties were used throughout these years and are typically replicated several times within a given trial.

Data was recored for four agronomic traits, grain yield (GY), plant height (PH), test weight (TW) and heading date (HD). GY and TW have no missing data, but 842 records are missing PH and 246 records are missing HD. All phenotypes were standardized by subtracting the mean and dividing by the standard deviation of each trait across all environments to preserve variance differences within environments (Supplementary Table A2).

The population was genotyped with genotyping by sequencing (Elshire et al. 2011, GBS) markers aligned to the International Wheat Genome Sequencing Consortium (IWGSC) Ref-Seq v1.0 wheat genome sequence of ‘Chinese Spring’ (IWGSC 2018, accepted, unpublished).

Markers were filtered for minor allele frequency of at least 0.01 (Figure A1), no more than 30% missing scores, and no more than 10% heterozygous calls. Missing marker scores were imputed using categorical random forest imputation by chromosome, and all heterozygous calls (< 2% of all calls) were subsequently replaced with the population mode (i.e. homozygous major allele). Marker scores are presented as boolean indicators of the minor allele. Further details of the CNLM dataset can be found in Appendix A1.

#### 3.1.2 W-GY population

The W-GY wheat dataset of 599 historical wheat lines from the CIMMYT Global Wheat Breeding program (Crossa et al. 2010) was used in this study due to its importance in genomic prediction of an inbred population with non-additive variation (Crossa et al. 2010; Martini et al. 2016). The W-GY dataset consists of genotypic values of all lines for grain yield in each of four environments. The genetic correlations between these environments ranged from −0.19 to 0.66 and can be found in Martini (2016). As performance between these environments is not highly correlated, we refer to grain yield performance in each environment as a trait. The dataset was used in its entirety with one exception. Of the 1,279 available DArT markers, only the 1,188 with known chromosomal positions as denoted by Crossa et al. (Crossa et al. 2010) were utilized in this study. This information was required to know which markers belonged to which subgenome, such that subgenome specific relationship matrices could be calculated.

### 3.2 Statistical framework

#### 3.2.1 Subgenome additive effects

To illustrate, we begin with a linear mixed model depicting environments as fixed effects and genotypes as random.

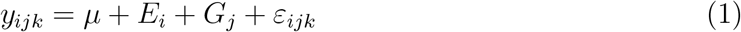

where *μ* is the population mean, *E_i_* and *G_j_* are the fixed environmental and random genetic effects, respectively, of the *j*^th^ genotype evaluated in the *i*^th^ environment, and *ε* is the error associated with the *k*^th^ observation. Using matrix notation, model (1) can be rewritten,

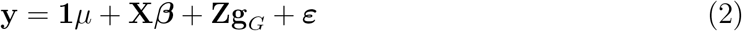

where **1**_*n*_ is a vector of ones, **X** is the design matrix, and ***β*** is the vector of fixed environmental effects. **Z** is the incidence matrix linking observations in the vector **y** to their respective genotype effects, in the vector **g**_*G*_. Normality was assumed for genotype effects and the residuals, where 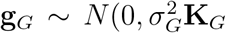 and 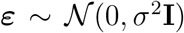. The genetic covariance, **K**_*G*_, can be derived from the expectation (or coefficient) of co-ancestry between individuals from a pedigree (Henderson 1985), or by an empirical estimation of the realized genetic relationship calculated with genome-wide markers (VanRaden 2008). When genome-wide markers are used to estimate *K*_*G*_, the genomic prediction model initially suggested by Nejati-Javaremi, Smith and Gibson (1997) and Meuwissen, Hayes and Goddard (2001) is obtained.

Given an *n* × *m* matrix, **M**, of *m* markers scored as reference allele counts (i.e. {0,1, 2}) for *n* individuals, method I of Van Raden (2008) finds the genetic relationship **K** as,

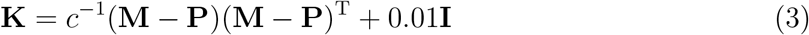

where **P** = **1**_*n*_ ⊗ 2**p**^*T*^ − 1, *c* = 2**p**^*T*^(**1** − **p**) and **p** is the vector of allele frequencies. The small coefficient of 0.01 was added to the diagonal to recover full rank after centering the matrix, such that **K**_*G*_ is invertible.

We use allohexaploid wheat to illustrate, but this method is easily truncated to allotetraploids, or extended to higher level allopolyploids. If we allow the total genetic effect, *G_j_*, to be decomposed into individual additive effects for each subgenome, such that *G_j_* = *A_j_* + *B_j_* + *D_j_*, the following model is obtained.

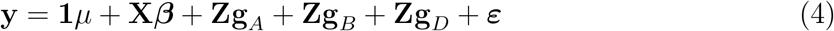

In model 4, each subgenome is allowed to have its own additive genetic variance and covariance between individuals, such that 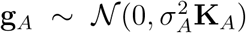, 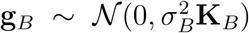 and 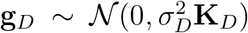. The realized additive genetic covariances for each subgenome, **K**_*A*_, **K**_*B*_ and **K**_*D*_, are estimated using only markers corresponding to the respective subgenome, and calculated as described above.

##### Subgenome epistatic interactions

Following Henderson (1985), the epistatic covariance of individuals can be calculated as the Hadamard product of the component covariance matrices. Martini et al.(2016) provide a proof of Henderson’s method using genome-wide markers to estimate the additive by additive covariance matrix, **H.** An additional linear kernel can then be added for an additive by additive epistatic interaction term, *I_j_*, once the additive covariance is estimated to obtain the following model.

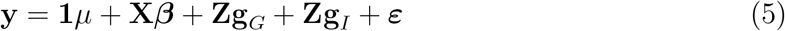

where 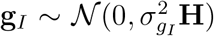. Martini (2016) does not report scaling **H** by the sum of the joint marker variances. Although this scalar does affect the parameter estimate, it does not not affect the model fit or prediction. We use the square of the sum of the marker variances as an approximation to avoid calculating all joint marker variances.

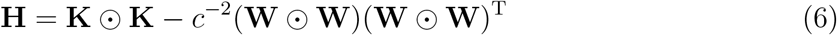

where **W** = **M** – **P**.

The additive by additive epistatic interaction term, **g**_*I*_, can also be decomposed into across subgenome interactions and within subgenome epistatic interactions such that 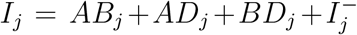 where *AB_j_, AD_j_* and *BD_j_* are the subgenome interaction effects and 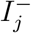 is the remaining epistatic effects due to within subgenome epistasis. Since no markers are shared across subgenomes, subgenome interaction covariances can be estimated by the Hadamard product of their component covariance matrices (Martini et al. 2016). These interactions can then be incorporated in the following model.

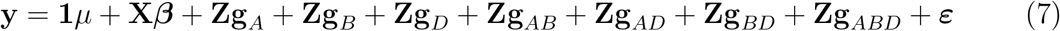

where 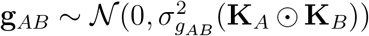, 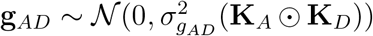, 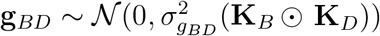 and 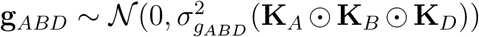. The three way interaction is included here for biological completeness, but was found to be estimated on the boundary (i.e. zero) for all traits, and was therefore dropped from further analyses.

### 3.3 Accounting for population structure

Under Hardy Weinberg equilibrium, subgenomes segregate independently, such that for subgenome effects, Cov(*A, B*) = Cov(*A, D*) = Cov(*B, D*) = 0 and Var(*G*) = Var(*A*) + Var(*B*) + Var(*D*). A breeding program, however, intentionally violates this assumption, and therefore may contain significant population structure. Price et al. (2006) demonstrated that the first *k* largest principal components (PCs) of the kinship matrix can be used to control for population structure in genome-wide association studies, and its use has since become wide spread. Because most realized estimates of additive covariance are proportional to **MM**^*T*^, singular value decomposition of **M**, instead of **MM**^*T*^, can be used to separate the population structure as the first few principal components from the entire matrix of marker scores before it is divided into its subgenome components. This is accomplished by first extracting the first *k* principal components in the *n* × *k* matrix **Q**. The marker matrix can then be reconstructed by setting the first *k* singular values of the diagonal matrix to zero and multiplying to produce a matrix 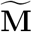 with the population structure removed from each subgenome simultaneously (Appendix A2).

Additive covariance matrices with reduced collinearity can then be calculated for each subgenome from 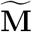 and incorporated into the model as previously described. **Q** can then be added to the model as a set of fixed covariates, with slopes ***γ***, such that the model will now be of the form

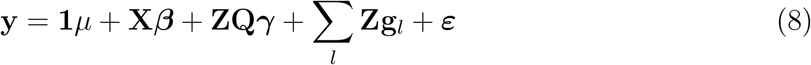

for all *l* genetic terms in the model. Genomic estimated breeding values are then predicted by summing the centered population structure and genetic effects. For this study, a population structure of dimension *k* = 5 was chosen for both the CNLM and W-GY datasets, and used to compare to the *k* = 0 models that do not correct for population structure.

### 3.4 Genomic prediction

To determine the predictability of genetic effects and the variability of variance component estimates, k-fold cross-validation with 5 folds was performed with 10 replications. For each replicate, the set of individuals was randomly split into five groups, with 4 groups of 289 and one of 291. For each fold, records of individuals in the fold were removed (i.e. masked) from the dataset. Each model was subsequently fit with the remaining lines and used to predict the whole genetic effect of the masked lines in the fold. Predictions for all five folds were gathered and correlated to the “true” genetic values once for each replicate. In this way, prediction results are directly comparable between the different models, and not subject to differences in the individuals sampled. The whole genome values were calculated as the sum of the genotypic additive and epistatic effects in the model as previously described. Due to the unbalanced nature of the CNLM dataset, “true” genetic values were calculated as in equation 2 but were considered independent with a covariance **K**_*G*_ = **I**.

### 3.5 Software

Models were fit using Restricted Maximum Likelihood (REML) with the software ASReml (Gilmour 1997) implemented in R (Butler 2009). The Tassel 5.0 GBS pipeline v2 (Glaubitz et al. 2014) along with the ‘bwa’ alignment tool (Li and Durbin 2009) were used for aligning GBS markers to the reference genome. All additional computation, analyses and figures were made using base R (R Core Team 2015) implemented in the Microsoft Open R environment 3.3.2 (Microsoft 2017).

### 3.6 Data availability

Phenotypes for the CNLM population are included in the file ‘pheno.txt’. Marker information and imputed marker scores for the CNLM population are included in files ‘snpInfo.txt’ and ‘snpMatrix.txt’, respectively. Best Linear Unbiased Predictors (BLUPs) for whole and subgenome additive effects (GEBVs and SGEBVS, respectively), as well as non-additive whole and subgenome interaction effects can be found in the ‘effectTable.txt’ file. Genotype and phenotype data for the W-GY population can be found in the ‘BGLR’ package of R (Campos and Pérez Rodriguez 2015), and marker chromosome information can be found in Crossa et al. (2010).

## 4 Results

### 4.1 Model fit and variance components

Model fit was assessed using Akaike’s Information Criterion (AIC). Whole genome models tended to have the lowest AIC values, with the exception of the PH and HD traits for the epistatic ABD×ABD models in the CNLM population. When whole genome models had lower AIC values, the comparable subgenome models had only marginally higher AIC values (Tables 1 and 2). Whole genome predictions between comparable whole genome and subgenome models were correlated at *ρ* > 0.999 or *ρ* > 0.993 for traits within the CNLM and W-GY populations, respectively. This indicates that little, if any, genetic information was lost by splitting the whole genome into biologically relevant subgenome effects. The lack of perfect correlation is at least partially due to floating point rounding errors during model fitting and summation of genotype effects.

**Table 1:**
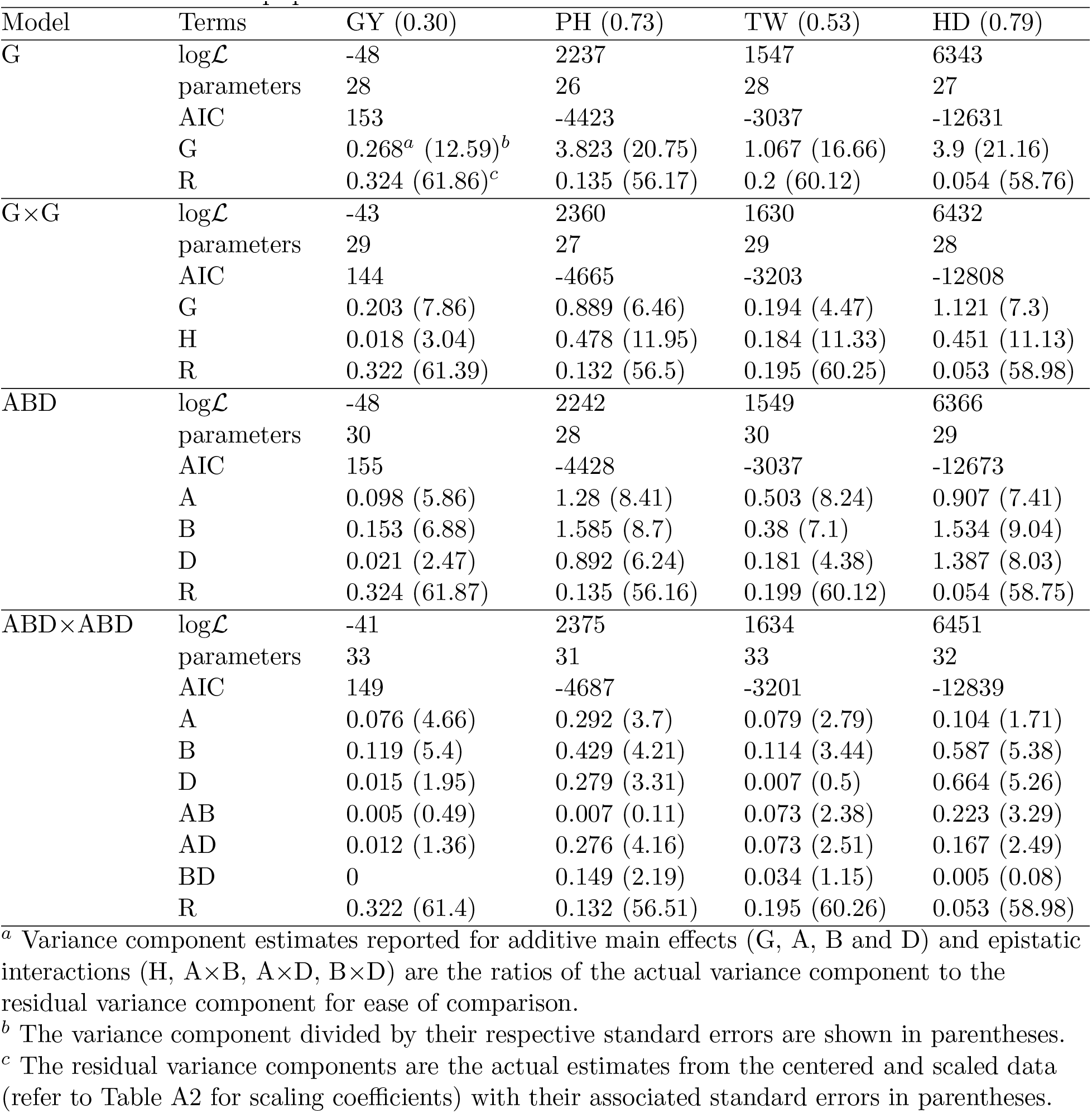
Table of model fit statistics for whole genome and subgenome genomic prediction models in the CNLM population.

**Table 2:**
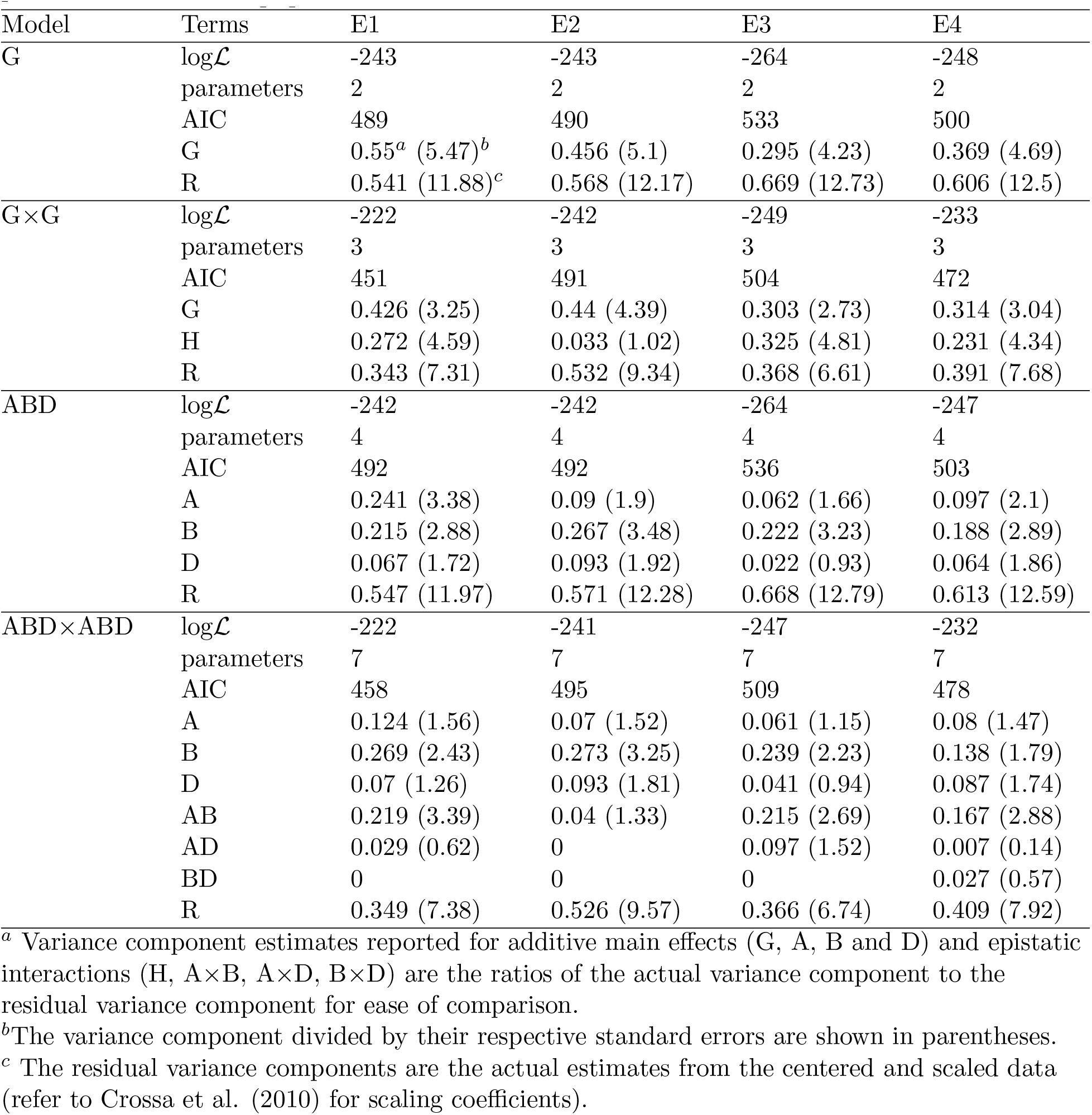
Table of model fit statistics for whole genome and subgenome genomic prediction models in the W-GY population.

| Model | Terms | E1 | E2 | E3 | E4 |
| --- | --- | --- | --- | --- | --- |
| G | $\log\mathcal{L}$ | -243 | -243 | -264 | -248 |
|  | parameters | 2 | 2 | 2 | 2 |
|  | AIC | 489 | 490 | 533 | 500 |
|  | G | 0.55<sup>a</sup> (5.47)<sup>b</sup> | 0.456 (5.1) | 0.295 (4.23) | 0.369 (4.69) |
|  | R | 0.541 (11.88)<sup>c</sup> | 0.568 (12.17) | 0.669 (12.73) | 0.606 (12.5) |
| G×G | $\log\mathcal{L}$ | -222 | -242 | -249 | -233 |
|  | parameters | 3 | 3 | 3 | 3 |
|  | AIC | 451 | 491 | 504 | 472 |
|  | G | 0.426 (3.25) | 0.44 (4.39) | 0.303 (2.73) | 0.314 (3.04) |
|  | H | 0.272 (4.59) | 0.033 (1.02) | 0.325 (4.81) | 0.231 (4.34) |
|  | R | 0.343 (7.31) | 0.532 (9.34) | 0.368 (6.61) | 0.391 (7.68) |
| ABD | $\log\mathcal{L}$ | -242 | -242 | -264 | -247 |
|  | parameters | 4 | 4 | 4 | 4 |
|  | AIC | 492 | 492 | 536 | 503 |
|  | A | 0.241 (3.38) | 0.09 (1.9) | 0.062 (1.66) | 0.097 (2.1) |
|  | B | 0.215 (2.88) | 0.267 (3.48) | 0.222 (3.23) | 0.188 (2.89) |
|  | D | 0.067 (1.72) | 0.093 (1.92) | 0.022 (0.93) | 0.064 (1.86) |
|  | R | 0.547 (11.97) | 0.571 (12.28) | 0.668 (12.79) | 0.613 (12.59) |
| ABD×ABD | $\log\mathcal{L}$ | -222 | -241 | -247 | -232 |
|  | parameters | 7 | 7 | 7 | 7 |
|  | AIC | 458 | 495 | 509 | 478 |
|  | A | 0.124 (1.56) | 0.07 (1.52) | 0.061 (1.15) | 0.08 (1.47) |
|  | B | 0.269 (2.43) | 0.273 (3.25) | 0.239 (2.23) | 0.138 (1.79) |
|  | D | 0.07 (1.26) | 0.093 (1.81) | 0.041 (0.94) | 0.087 (1.74) |
|  | AB | 0.219 (3.39) | 0.04 (1.33) | 0.215 (2.69) | 0.167 (2.88) |
|  | AD | 0.029 (0.62) | 0 | 0.097 (1.52) | 0.007 (0.14) |
|  | BD | 0 | 0 | 0 | 0.027 (0.57) |
|  | R | 0.349 (7.38) | 0.526 (9.57) | 0.366 (6.74) | 0.409 (7.92) |

<sup>a</sup> Variance component estimates reported for additive main effects (G, A, B and D) and epistatic interactions (H, A×B, A×D, B×D) are the ratios of the actual variance component to the residual variance component for ease of comparison.
<sup>b</sup>The variance component divided by their respective standard errors are shown in parentheses.
<sup>c</sup> The residual variance components are the actual estimates from the centered and scaled data (refer to Crossa et al. (2010) for scaling coefficients).

Subgenome additive variance parameter estimates were positive for all models, but subgenome interaction variance parameter estimates were often estimated on the boundary (i.e. near zero). Variance parameters estimated on the boundary were thus considered to be exactly zero. Shifts in variance component importance were seen when the epistatic terms were added in the model. For example, for the TW and E1 traits in the CNLM and W-GY populations, respectively, the A genome component was the largest in the additive only model, but was reduced to less than that of the B genome component in the epistatic model. Additive variance components were generally reduced in epistatic models compared to additive only models, but this reduction in additive variance was accompanied by non-zero subgenome interaction components. The B genome contributed the greatest amount of additive variance in the epistatic ABD×ABD models for all traits accept HD. While the D genome variance component was far smaller than the A subgenome component for GY and TW in the CNLM population, it was comparable to the A subgenome component for all traits in the W-GY population.

The A×B component was particularly important for the W-GY traits, E1, E3 and E4, as well as the HD and TW in the CNLM population. The A× D component also featured prominently for the PH and TW traits in the CNLM population. The B× D component appeared to be less important, having the largest effect for PH. No epistatic terms were significantly greater than zero for the E2 trait in the W-GY population. Addition of epistatic interactions resulted in a significant likelihood ratio test at p < 10^−6^ for all traits except GY, which was significant at p < 10^−2^. Despite the significant addition of epistatic terms, additive only GEBVs were highly correlated with whole genome predictions from the epistatic models, at *ρ* ≥ 0.988 for the CNLM population and *ρ* ≥ 0.869 for the W-GY population. A model containing the three-way subgenome epistatic term was fit for all traits, but estimates of the three-way interaction variance parameter were zero for all traits.

The distributions of variance component estimates from repeated sub-sampling of the data during *k*-fold cross-validation were centered near the point estimate from the full model fits. These distributions were either as wide (≈ 2 standard errors from the center) or tighter than expected based on the standard error from the full model fit (Figures 1 and 2). Standard errors were generally larger for epistatic variance components relative to their magnitude than additive variance components. Standard errors relative to their respective parameter estimates tended to be larger for all terms in models with more estimated variance parameters (Tables 1 and 2).

**Figure 1:**
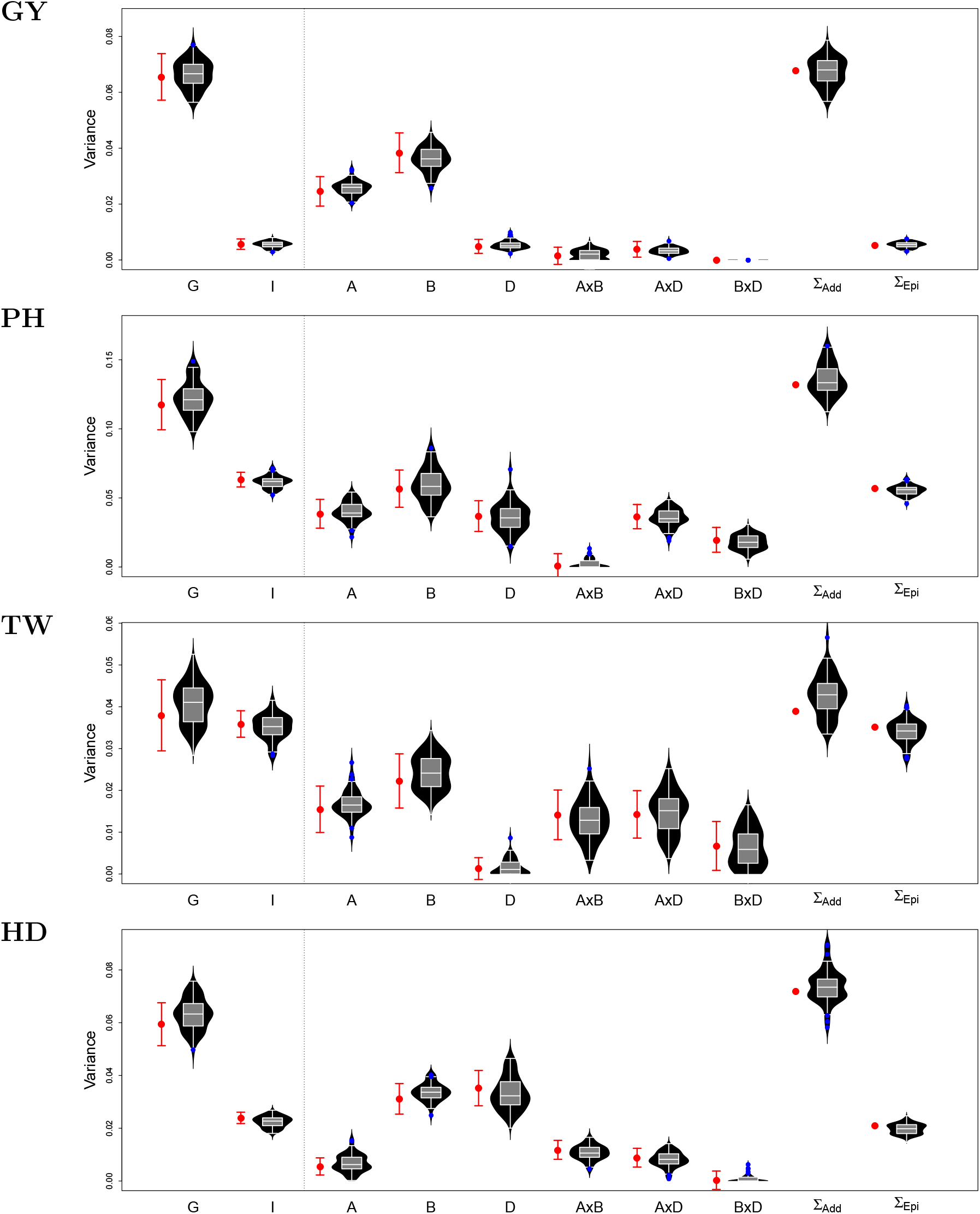
Estimates and standard errors of variance components for four traits in the CNLM populations from the full model (red) compared to the sampling distribution of variance component estimates from the cross-validation scheme (black violins). G× G and ABD× ABD models are shown to the left and right of the dotted line, respectively. The sum of the additive and interaction variance components is also shown for the ABD×ABD model.

**Figure 2:**
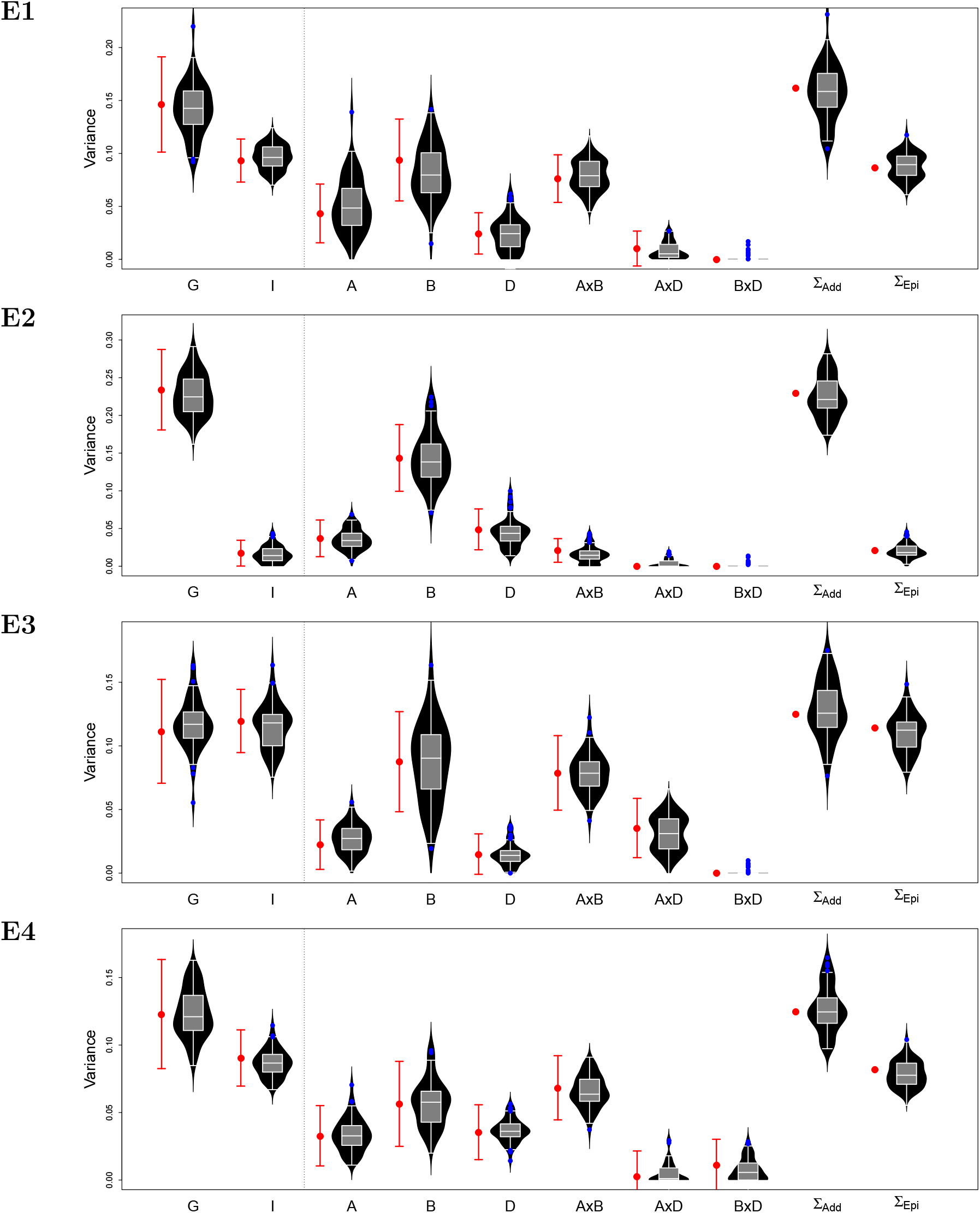
Estimates and standard errors of variance components for four traits in the W-GY populations from the full model (red) compared to the sampling distribution of variance component estimates from the cross-validation scheme (black violins). G× G and ABD× ABD models are shown to the left and right of the dotted line, respectively. The sum of the additive and interaction variance components is also shown for the ABD × ABD model.

### 4.2 Subgenome additive effects

Subgenome estimated breeding values (SGEBVs) were moderately correlated with the whole genome effect, but weakly correlated with one another (Tables 3 and 4). The individuals with the highest SGEBV for one subgenome never had the highest SGEBV for the other two subgenomes, and were often not in the top 95% quantile of the population based on the other two subgenomes (Figures 3 and 4). For example, the individuals with the highest A, B and D SGEBV for GY in the CNLM population ranked 43^rd^, 39^th^ and 60^th^ for the whole genome effect, respectively. In contrast, the individual with the best A SGEBV for GY ranked 1067^th^, 952^nd^ for the B and D genome, respectively. The individual with the highest B genome breeding value for GY ranked 221^th^ and 1393^rd^ for the A and D genomes, respectively. The individual with the highest D genome breeding value for GY ranked 347^th^ and 123^rd^ for the A and B subgenome, respectively. The individual with the highest whole genome GEBV for grain yield ranked 6^th^, 22^nd^ and 519^th^ for the A, B and D SGEBV, respectively. In several cases, the top individual based on a SGEBV was not in the top 95% quantile based on their whole genome effect, particularly in the W-GY population.

**Figure 3:**
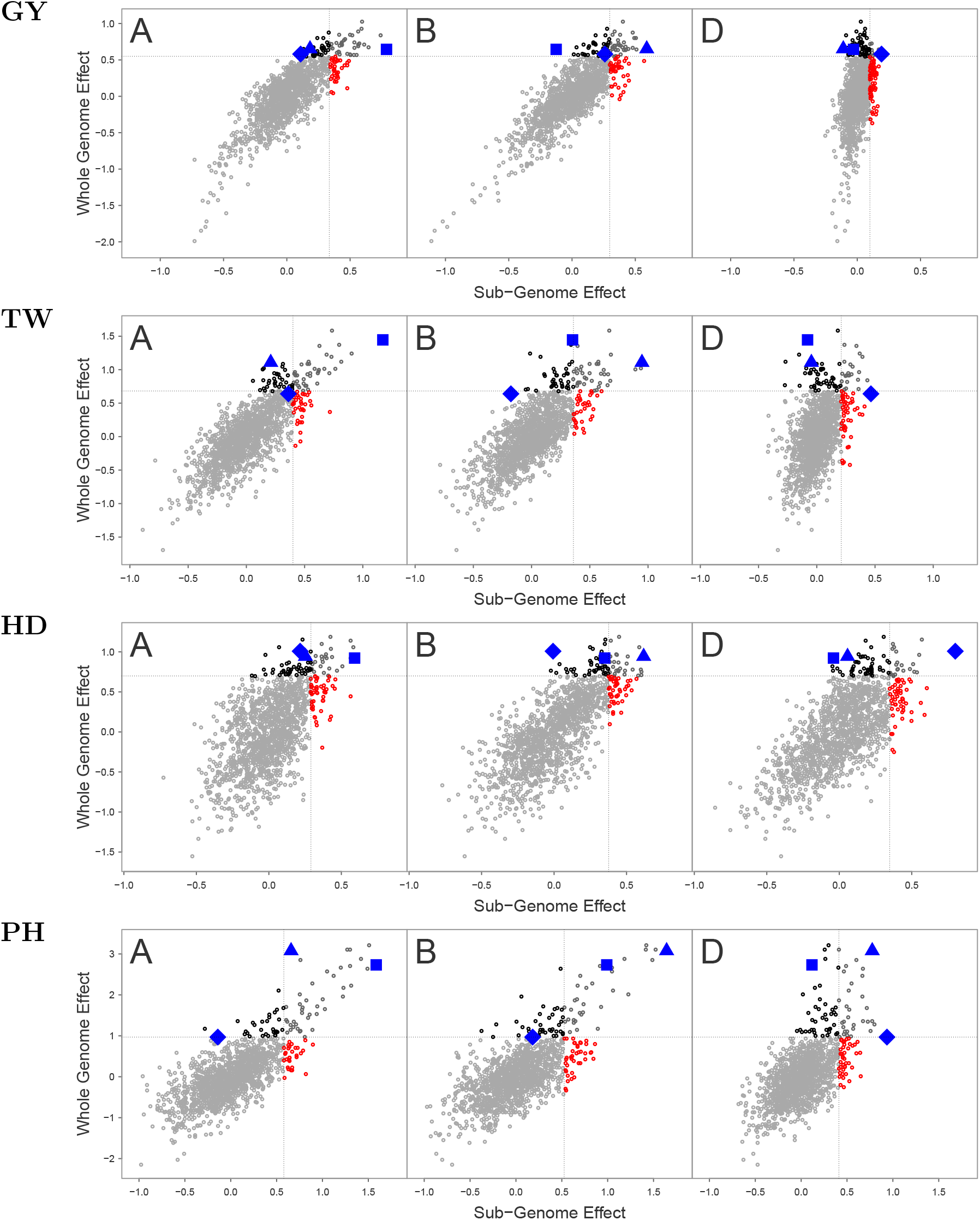
Plot of whole genome additive effects (GEBV) by subgenome additive effects (SGEBV) for four traits in the CNLM populations. The dotted line indicates the 95% quantiles for whole or subgenome effects. Blue squares, triangles and diamonds indicate the line with the highest SGEBV for each of the A, B and D subgenomes, respectively.

**Figure 4:**
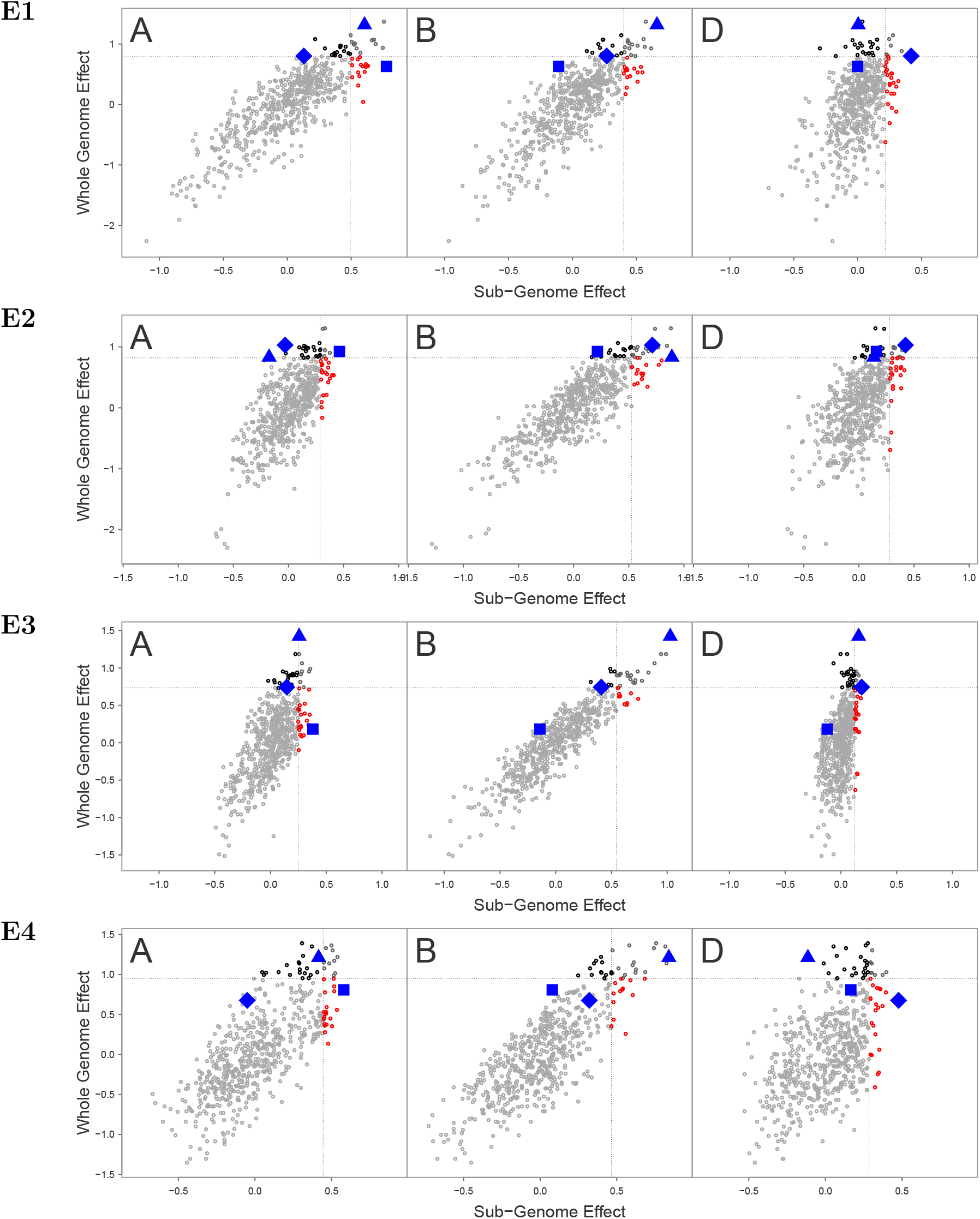
Plot of whole genome additive effects (GEBV) by subgenome additive effects (SGEBV) for four traits in the W-GY populations. The dotted line indicates the 95% quantiles for whole or subgenome effects. Blue squares, triangles and diamonds indicate the line with the highest SGEBV for each of the A, B and D subgenomes, respectively.

**Table 3:**
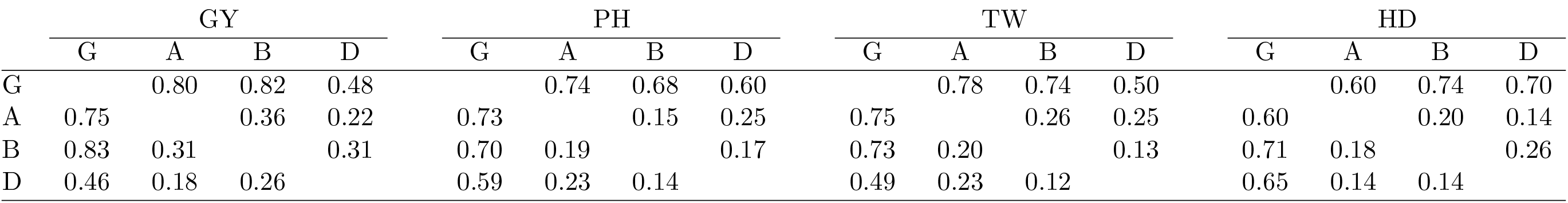
Correlation of whole genome and subgenome additive effects in the CNLM population. Correlations of additive random effects without correcting for population structure are shown above the diagonal, while correlations of effects correcting for populations structure using the first *k* = 5 PCs is shown below the diagonal.

**Table 4:**
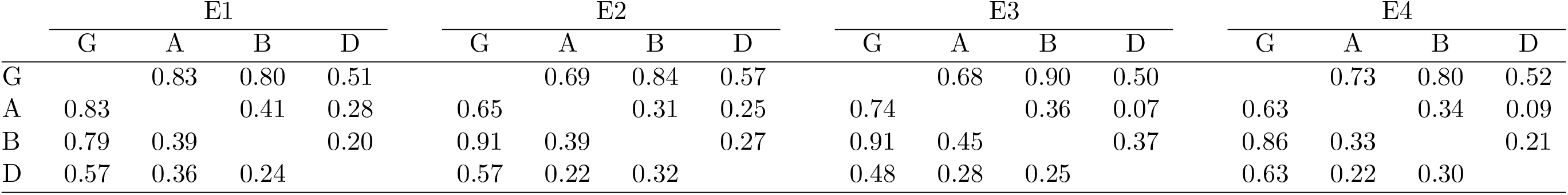
Correlation of whole genome and subgenome additive effects in the W-GY population. Correlations of additive random effects without correcting for population structure are shown above the diagonal, while correlations of effects correcting for populations structure using the first *k* = 5 PCs is shown below the diagonal.

### 4.3 Prediction accuracy

Including epistasis kernels significantly improved genomic prediction accuracy for all traits except GY and E2 (Table 5). Subgenome models had either comparable or slightly lower mean prediction accuracy than whole genome models for all traits except HD, for which subgenome models had superior accuracy. The variability in the prediction accuracy based on the individuals sampled was either the same (GY and TW) or lower (PH and HD) for the epistatic models compared to the additive models in the CNLM population, but was similar in the W-GY population (Table 5). The variability in prediction accuracy was increased for the subgenome models compared to the whole genome models in the W-GY population for some traits (E2 and E3), but was either the same or decreased in the the CNLM population.

**Table 5:** Table of genomic prediction accuracies for eight traits in the CNLM (GY, PH, TW and HD) or W-GY (E1, E2, E3, E4) populations with *k* = 0 and *k* = 5.

| CNLM | $k$ | GY | PH | TW | HD |
| --- | --- | --- | --- | --- | --- |
| G | 0 | 0.601<sup>a</sup> (0.008)<sup>b</sup> | 0.559 (0.007) | 0.515 (0.010) | 0.664 (0.009) |
| ABD |  | 0.600 (0.008) | 0.557 (0.008) | 0.514 (0.011) | 0.679 (0.007) |
| GH |  | 0.604 (0.008) | 0.637 (0.004) | 0.576 (0.010) | 0.712 (0.008) |
| ABD×ABD |  | 0.603 (0.008) | 0.638 (0.005) | 0.569 (0.011) | 0.720 (0.006) |
| G | 5 | 0.600 (0.009) | 0.558 (0.007) | 0.514 (0.011) | 0.663 (0.010) |
| ABD |  | 0.600 (0.009) | 0.556 (0.008) | 0.513 (0.011) | 0.678 (0.008) |
| GH |  | 0.602 (0.008) | 0.624 (0.005) | 0.560 (0.010) | 0.701 (0.008) |
| ABD×ABD |  | 0.602 (0.007) | 0.618 (0.005) | 0.557 (0.010) | 0.708 (0.006) |
| W-GY | $k$ | E1 | E2 | E3 | E4 |
| G | 0 | 0.501 (0.010) | 0.493 (0.016) | 0.356 (0.008) | 0.457 (0.010) |
| ABD |  | 0.492 (0.012) | 0.481 (0.023) | 0.346 (0.010) | 0.449 (0.011) |
| GH |  | 0.568 (0.010) | 0.494 (0.017) | 0.396 (0.013) | 0.520 (0.010) |
| ABD×ABD |  | 0.549 (0.011) | 0.484 (0.023) | 0.393 (0.015) | 0.509 (0.013) |
| G | 5 | 0.502 (0.010) | 0.491 (0.017) | 0.354 (0.007) | 0.458 (0.010) |
| ABD |  | 0.495 (0.011) | 0.475 (0.024) | 0.345 (0.010) | 0.453 (0.011) |
| GH |  | 0.526 (0.010) | 0.491 (0.017) | 0.381 (0.007) | 0.493 (0.011) |
| ABD×ABD |  | 0.520 (0.012) | 0.475 (0.023) | 0.373 (0.013) | 0.486 (0.012) |

<sup>a</sup> Mean genomic prediction accuracy across ten replicates of five fold cross validation.
<sup>b</sup> Standard deviation of prediction accuracy across ten replicates are shown in parentheses.

### 4.4 Adjustment for population structure

The first two principal components explained 17% and 19% of the variance of M in the CNLM and W-GY populations, respectively, indicating that some population structure exists in both populations (Supplementary Figure S1). The correlation of additive genetic covariance estimates between individuals based on the three subgenomes declined as PCs were removed from **M**, but appeared to level out between 5 to 10 PCs (Figure 5). Correlation of whole genetic effects between additive models, G and ABD, for *k* = 0 and *k* = 5 was ≥ 0.999 and ≥ 0.996 for the CNLM and W-GY populations respectively. Whole genome effect correlations were lower between epistatic models G×G and ABD×ABD, with coefficients of ≥ 0.998 in the CNLM population and ≥ 0.980 in the W-GY population.

**Figure 5:**
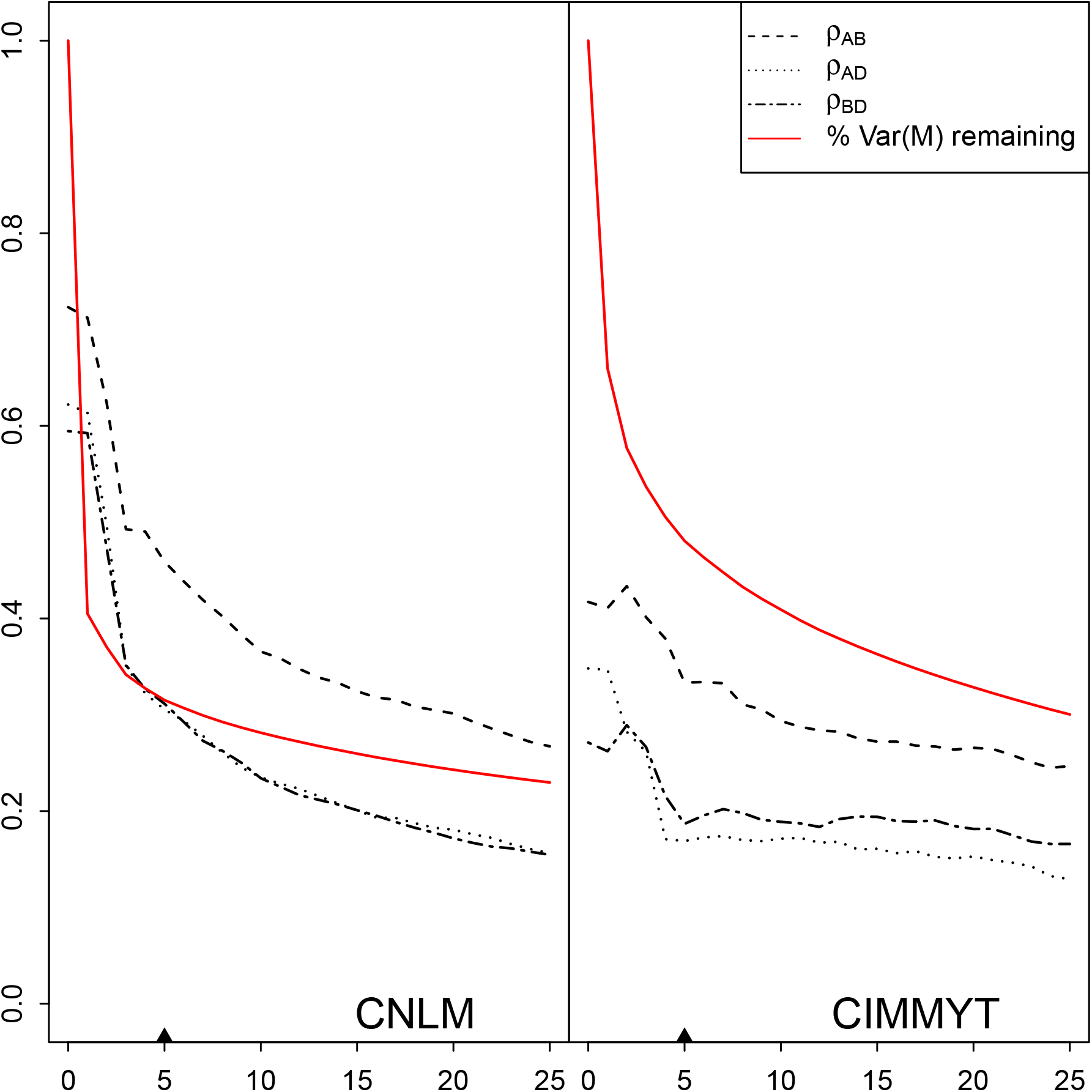
Correlation coefficient, *ρ*, of off-diagonal elements of estimated additive covariance matrices **K**_*A*_, **K**_*B*_ and **K**_*D*_. The percent genotype marker variance remaining in the marker matrix after removing *k* dimensions is shown in red. The chosen population structure dimension *k* = 5, is indicated by a ▲.

Removing population structure with *k* = 5 reduced most of the SGEBV effect correlation coefficients by up to 0.06 in the CNLM population, but there was one instance in which one correlation coefficient increased from 0.14 to 0.19 between A and B SGEBVs for PH (Table 5). This was not the case for the W-GY population, where many of the SGEBV effect correlations increased by up to 0.21.

Variance components generally decreased as *k* was increased from 0 to 10 (Figure 6). Ranks of additive variance components relative to one another were stable for most traits, while epistatic variance components were more sensitive to changes in *k*. Significant epistatic variance component rank changes occurred for the PH, TW and E4 traits. For PH, the A× D term was comparable in magnitude with the additive variance components for A and D when *k* = 0, but declined as *k* increased. The reduction in A× D variance for PH was accompanied by an increase in both the A× B and B× D terms. Similarly, a decline in A× B variance was followed by an increase in B× D for TW and A× D for E4 as *k* was increased.

**Figure 6:**
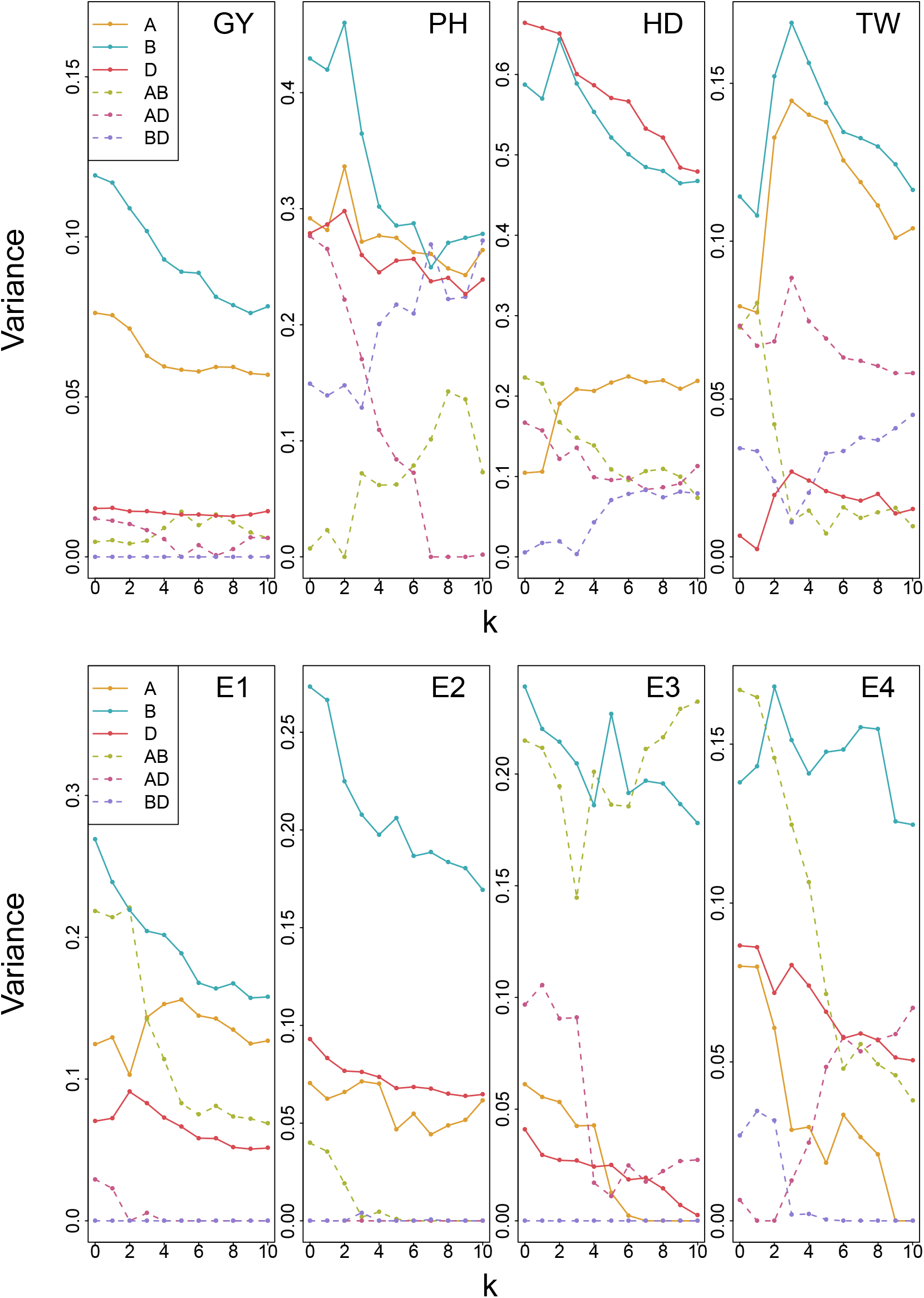
Subgenome additive and interaction variance parameter estimates from the ABD×ABD model correcting for population structure with *k* ∈ {0,1,…, 10} principal components as fixed effects. Models were fit with four traits for the CNLM population and four traits for the W-GY population.

Correlations of variance component estimates were calculated from the average information matrix for models *k* ∈ {0,1,…, 10} (Supplementary Figures S2 and S3). Correlations between subgenome additive variance estimates were generally low (0.2-0.4), while correlations of subgenome interaction variance estimates were high (0.8-0.95), and correlations between the two were moderate (0.4-0.6). Despite a small reduction in the correlation of SGEBVs as *k* was increased from 0 to 5, little reduction in variance component estimate correlations was observed as *k* was increased from 0. Generally, correlations of additive variance parameter estimates were slightly reduced while correlations between interaction variance parameter estimates increased slightly.

## 5 Discussion

### 5.1 Model fit and variance components

While whole genome models tended to be the most parsimonious, subgenome models are worth consideration because they provide insight into the biology of the allopolyploid organism. Given the stability of variance component estimation and that no genetic information appears to be lost by partitioning the whole genome into its individual subgenome additive effects, such a partition is informative.

The method presented here could be used for any set of independent loci, such as estimating a variance component and breeding value for each chromosome. However, this will become computationally burdensome as the number of variance components to be estimated increases. If the number of variance parameters to estimate is high and the data set is small this may become infeasible. It is also unclear if the estimates from larger numbers of additive kernels would be reliable.

Bernardo and Thompson (2016) assigned a breeding value for each of the 10 maize chromosomes by fitting a single ridge regression model to estimate marker effects. They subsequently summed marker effects by chromosome to produce a breeding value for each chromosome. However, this method does not allow for direct estimation of variance components for each unit of chromatin. By fitting each unit simultaneously, variance attributable to sets of loci will be split, and the sum of the variance estimates should not exceed the total genetic variance. It is unclear what effect LD across chromosomes has on the variance parameters estimated.

Here we assumed that the subgenome effects are independent, but this is clearly not the case. Generally, we can express the genetic variance due to the three sub genomes as Var (vec([**g**_*A*_ **g**_*B*_ **g**_*D*_])) = **S** ⊗ **J**_*n*_ ⊙ **K**, where **S** is the subgenome covariance matrix, **J** is an *n* × *n* matrix of ones for *n* genotypes, and **K** is the additive relationship matrix for within and across subgenomes. In this report, we have assumed that **S** is diagonal with 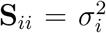 for the *i*^th^ subgenome, and **K** is a block diagonal with the *i*^th^ diagonal block represented by the subgenome additive covariance matrix.

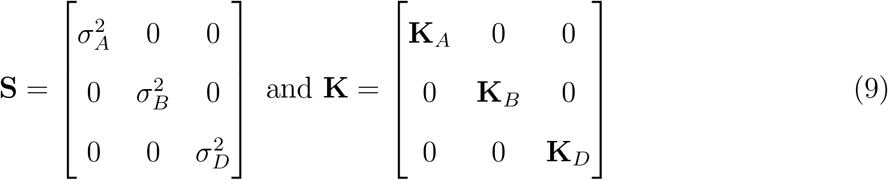

An unstructured covariance matrix, **S**, could be estimated, with correlation coefficients between subgenomes. The subgenome effects would be allowed to have a correlation such that

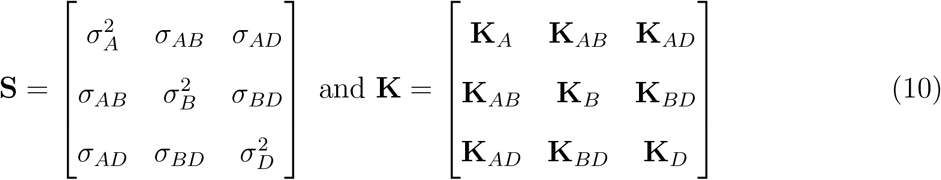

However, it is unclear what the covariance structure should be between subgenomes (e.g. **K**_*AB*_). If consensus haplotypes from uniquely identifiable sequences could be determined with two or more alleles segregating in at least two subgenomes, a covariance across the subgenomes could be constructed. Polymorphisms that predate speciation would be used to identify the consensus haplotypes, while post speciation polymorphisms would be used to identity the subgenome origin. Individuals would then receive a score based on the number of consensus haplotypes they have in common between two subgenomes. This could prove to be a formidable challenge given the evolutionary time between the subgenome ancestors.

The Hadamard product of the two additive covariance matrices is a tempting candidate for these off diagonal blocks, however, this would substitute a correlation coefficient between additive effects in place of an epistatic variance. It is unclear to these authors if epistasis variance can be thought of and modeled as a correlation between additive effects.

### 5.2 Genetic architecture

The genetic architecture of grain yield (GY, E1, E2, E3, E4) in the two populations investigated here are markedly similar, despite the divergent genetic backgrounds of the two populations. The CNLM population is primarily comprised of breeding lines and varieties derived from germplasm historically grown in the North East, in contrast to the W-GY population which has a broader pedigree.

The D genome is known to have low genetic diversity due to limited gene flow from a single *Ae. tauschii* lineage after the most recent allopolyploidization event (Wang et al. 2013), estimated to have taken place as recently as 10,000 years ago (Salamini et al. 2002; Marcussen et al. 2014). The International Maize and Wheat Improvement Center (Centro Internacional de Mejoramiento de Maíz y Trigo, CIMMYT) has introgressed some genetic material from the D genome ancestor, *Ae. tauschii*, through the use of synthetically produced hexaploid wheat to increase the genetic diversity of the historically bottle-necked D genome. The higher proportion of D genome variance in the W-GY population may be due to the increased use of wild *Ae. tauschii* in their breeding program, highlighting the merit of the strategy.

Many of the subgenome epistatic variance parameters were estimated at zero, possibly due to a lack of power to detect them. Greater genetic diversity, larger numbers of individuals, and higher allele frequencies would allow for increased power to detect true interactions. Hill et al. (2008) emphasized the effect of low allele frequencies on epistatic interactions, proving that as allele frequencies (and therefore joint frequencies of alleles at two loci) approach zero or one, most of the epistatic variance becomes additive. For example, suppose two loci have a large interaction, such that one pair of alleles is selected. Once one locus becomes fixed, all remaining variance is due to seggregation at the other locus, and becomes strictly additive. The low joint frequency is magnified in the three way interactions, likely causing the inability to detect any three way epistatic interaction signal between the three subgenomes.

This is also apparent in the reduction of additive variance components upon the addition of epistatic terms to the model. These components were often estimated to be rather large compared to the additive components, but did not change the final whole genome value drastically. This suggests that the additive terms absorb much of the epistatic variance in the absence of the epistatic kernels.

The A×B epistatic terms were the most important for many of the traits, reflecting the greater genetic variation of these two subgenomes. Subgenome interaction terms including the D genome were notably more important for traits known to have important loci on the D genome. PH is partially governed by two dwarfing genes, *Rht-1D* and *Rht-1B* on 4B and 4D, respectively. These two genes have been shown to exhibit a less than additive epistatic interaction, where the double wildtype is less tall than expected based on the additive effects of the two semi dwarfs from the double dwarf. The B× D term was large for PH, particularly after correction for population structure. Population structure is common for these genes, as breeding programs primarily utilize one or the other dwarfing gene to avoid producing double dwarfs during crossing, which are agronomically undesirable.

### 5.3 Selection on SGEBVs

Partitioning genetic variance to the subgenomes of an allopolyploid provides a method for identifying individuals with complementary subgenomes as potential parents for crossing. If we consider the upper 95% quantiles as candidates for parental selection, many of the top candidates based on subgenome breeding values would not be considered candidates based on their whole genome breeding value. When they would be considered, they were typically not the top candidates. The low correlation between SGEBVs highlights the opportunity to identify individuals with complementary subgenomes for crossing. These individuals may or may not be among the top performing selection candidates, demonstrating that the optimum set of crosses are not always between the top performing individuals (Akdemir and Sánchez 2016).

The low correlation and high predictability of SGEBVs suggests that individual subgenomes may be directly manipulated as never before. Prior to the discovery and use of genetic markers to track genomic regions, the phenotype (or some summary statistic thereof) was the only indicator of the genetic structure of a genetically distinct individual. Variety releases still demonstrate this legacy, with phenotypic descriptors that define a new variety as genetically distinct from other similar varieties. One breeding strategy will be selecting parents for crossing that have complementary SGEBVs to increase the potential of transgressive segregation in the resulting offspring. We envision other breeding strategies beyond simply choosing parents with complementary subgenomes, and see an opportunity to weight SGEBVs according to some breeding goal.

For example, a newly formed population could undergo several rounds of genomic selection only on the D genome SGEBVs (i.e. weights of 0, 0 and 1 for the A, B and D subgenomes respectively) before phenotypic or whole genome selection. Because the D genome contributes the least to the total genetic variance, phenotypic selection on D genome loci is challenging. Selection will act on the largest sources of genetic variance first, potentially leading to fixation of small effect loci in the D genome by drift, while selection acts on the large effect loci on the A and B genomes first. By selecting on D genome SGEBVs, gains can be made to the D genome directly with little to no selection on the A and B genomes, a feat previously impossible with phenotypic selection.

### 5.4 Subgenome interactions

Genomic prediction of GY and E2 did not appear to benefit from including epistatic interactions as it did for the other six traits. This may be due in part to the highly polygenic nature of grain yield, which is the culmination of essentially all functional genetic variants subjected to stress throughout the growth cycle. The E2 trait in the W-GY population has previously been shown to be invariant to the addition of various epistatic terms (Crossa et al. 2010; Martini et al. 2016), and it is unclear why this population does not exhibit non-additive variation in this environment. It may be that important epistatic interactions of GY in the CNLM population are too small to detect or are involved with differing performance across years or locations, such that they are lacking in a model that does not include genotype by environment interactions.

Subgenome epistatic terms increased genomic prediction accuracy equivalent to modeling all pairwise interactions across the subgenomes, suggesting that the most important interacting loci are on different subgenomes. This result is consistent with the observation that newly formed allopolyploids undergo considerable changes in gene expression, known as genome shock (McClintock 1984). This shock has been suggested to be caused by incompatibilities of genetic pathways across the subgenomes (Comai et al. 2003). Residual subgenome incompatibility may still be affecting the germplasm pool, even thousands of years after the last polyploidization event. Decay of negative gene interactions has been shown to take hundreds to thousands of generations before all interacting genes are lost or silenced (Lynch and Conery 2003).

It is unclear what proportion of this non-additive signal is due to homeoallelic interactions. The proposed method models all pairwise interactions across subgenomes, of which homeoallelic gene interactions are a small minority in number. Smaller homeoallelic regions or homeoallele specific marker sets would need to be constructed to determine the relative importance of these interactions relative to other gene interactions across the subgenomes. The usefulness of the epistatic subgenome interactions is currently unclear and warrants further investigation.

Regardless of the source of the epistasis, we suggest that a breeding scheme should be designed to take advantage of beneficial subgenome interactions. If a suitable training set related to the breeding material can be established, subgenome interactions can be predicted in new, genotyped breeding materials. We suggest that a series of small bi-parental populations be constructed from important contributors to the breeding program, and be used in the development of a training population to balance high genetic diversity and high allele frequencies. This training population will be used to predict SGBEVs and subgenome interactions in individuals formed from new crosses. Individuals that contain favorable interactions can then be selected such that they are fixed in early filial generations. After fixation in a given line, phenotypic, whole genome, or subgenome selection can be used for further line development until complete homozygosity is reached.

### 5.5 Adjustment for population structure

The efficacy of the proposed method to handle population structure may need to be improved, or a different approach may need to be taken. While this method reduced the correlation of additive genetic subgenome covariance estimates across the three genomes, variance parameter estimate correlations were not drastically reduced. The correlation of subgenome interaction variance parameter estimates tended to increase slightly when accounting for higher levels of population structure, counter to the assumption that removing this structure should result in better estimates of subgenome interactions.

The lower correlation between epistatic models that correct and do not correct for population structure is likely due to removing Q from the marker matrix. Correcting for population structure also had a small, but negative effect on genomic prediction accuracy for epistatic models. The population structure fixed effect predictors are strictly additive and the loss of accuracy may be due to epistasis variance associated with these PCs (i.e. population structure epistasis). Epistatic variance related to these PCs may be recovered by using the squares of the PC scores, although this was not done in this study. At least for the additive models, it appears little to no genetic information is lost using the population structure adjustment proposed here.

Determining the best value for *k* will be at the crux for implementing this methodology for various traits and populations. The same population may need different values of *k* for different traits, depending on how the population is structured. Traits such as PH or HD may have less complex structure than traits such as TW or GY, due to different marker effect distributions and the history of the breeding population. Several methods might be used to determine *k* empirically from the marker matrix (Patterson, Price, and Reich 2006, e.g.), however, these methods may not capture subtle differences in the population structure of a given trait. We used the first one to *k* PCs in this study, but there is no reason why we must include all PCs up to some value *k*. There may be certain PCs that are important for a given trait, and could be tested as fixed effects for inclusion or exclusion.

This method may have better performance in populations with greater degrees of population structure than in the populations presented here. Use of this method for partitioning genetic variance to biologically important sets of chromatin and estimating epistatic interactions will need further testing and validation before widespread use.

## 6 Conclusion

To our knowledge, we provide the first attempt to assign a breeding value to each subgenome of an allopolyploid crop. With estimates of subgenome additive effects, parents with complementary subgenomes can be selected for crossing. Weighted selection of subgenomes using genomic prediction could be key to increasing the diversity of the D genome in wheat germplasm. Direct selection on the D genome may allow targeted introgression from *Ae. tauschii* while mitigating the effects of introducing unimproved alleles. Subgenome additive genetic variances appear to be estimated well, and no genetic information is lost partitioning the genome into its subgenome components. This demonstrates that partitioning genetic variance to the subgenomes of an allopolyploid can provide useful information for genomics assisted breeding efforts.

Subgenome interactions increase prediction accuracy, but it is unclear how well the epistatic variance is partitioned to the three interaction terms and what proportion of that variance is due to homeologous gene interactions. Because the homeologous interactions make up relatively few of the possible interactions across subgenomes, they may only explain a small portion of the observed epistatic variance. Yet, seeing as how homeologous genes likely operate in the same or similar physiological pathway, the likelihood for interactions between homeologous loci is high. Further research is needed to investigate the efficacy of modeling subgenome interaction terms, and to what degree this is explained by interactions between homeologous orthologs.

Allopolyploids have traditionally been treated as diploids in breeding programs because they undergo disomic inheritance. With modern DNA marker technology and ever increasing computational power, breeders of allopolyploids can further exploit the genetic complexity of their crops.

## 7 Acknowledgments

Funding of this research was provided by the USDA National Needs Fellowship for N. Santantonio, in partial fulfillment of the requirements for a Ph.D in Plant Breeding and Genetics at Cornell University. Additionally, the field trials comprising the phenotypic data for the CNLM population were funded in part by the Hatch Project # 149-447. Genotyping was funded by the Wheat Coordinated Agricultural Project (WheatCAP). We are also grateful to Jesse Poland’s research group at Kansas State University for their contribution to genotyping of CNLM materials. The authors thank the International Wheat Genome Sequencing Consortium for pre-publication access to IWGSC RefSeq v1.0. Finally, we would like to acknowledge the Cornell small grains staff, particularly David Benscher and James Tanaka, who were vital in implementing, collecting and processing the materials used to build the CNLM dataset.

## A1 Appendix 1 CNLM dataset

**Table A1:** Number of phenotypic observations for each location across 10 years.

| Year | Helfer | Ketola | McGowan | Snyder |
| --- | --- | --- | --- | --- |
| 2007 | 0 | 0 | 246 | 0 |
| 2008 | 0 | 432 | 0 | 426 |
| 2009 | 409 | 0 | 431 | 0 |
| 2010 | 311 | 304 | 0 | 305 |
| 2011 | 310 | 319 | 318 | 0 |
| 2012 | 307 | 306 | 0 | 302 |
| 2013 | 127 | 130 | 338 | 0 |
| 2014 | 232 | 232 | 0 | 233 |
| 2015 | 469 | 464 | 461 | 0 |
| 2016 | 427 | 425 | 0 | 428 |

Field plots 1.5 m by 3 m in size were planted with 100 g of seed in September or October of the year prior to the harvest year. Data was recorded for four agronomic traits: grain yield (GY), plant height (PH), test weight (TW) and heading date (HD). Plots were harvested for grain with a plot combine after physiological maturity and oven dried to a grain moisture of approximately 12%. Dried grain was cleaned, weighed and measured for moisture content using a grain moisture analyzer (GAC 2100, Dickey-John). GY was standardized to a uniform grain moisture of 12%. PH was measured as the distance from the ground to the top of the grain head at full extension. TW is used as a measure of grain quality and was measured as the mass of a volume of grain (*gL*^−1^) using the grain moisture analyzer (GAC 2100, Dickey-John) which corrects TW for moisture content. HD is a proxy for flowering time and was defined as the number of Julian days until 50% of the primary grain heads have extended out of the boot.

The data set initially consisted of 1,552 lines evaluated in 10,069 in 1.5 m by 3 m plots planted in September to October of the year prior to the harvest year. Thirty one lines from 2007 were not harvested for GY, nor were they genotyped, and were dropped from the data set. Because GY is of primary interest for breeding, plots that were not harvested or had missing values for GY were dropped, resulting in 9,090 plots with GY measurements.

**Table A2:** Means (*μ*) and standard deviations (*σ*) of four traits in the CNLM population.

| | units | $\mu$ | $\sigma$ |
| --- | --- | --- | --- |
| GY | kg ha<sup>-1</sup> | 5315.20 | 1015.76 |
| PH | cm | 90.84 | 11.99 |
| HD | Julian days | 151.64 | 3.87 |
| TW | g L<sup>-1</sup> | 74.95 | 3.09 |

This also caused two additional lines with missing GY measurements to be dropped from the dataset.

Due to the reasonable size of the dataset, small physical area of most trials, lack of replication within environment for most lines and the availability of genetic markers, raw plot observations were used and no attempt was made to correct plot level data for various spatial effects or otherwise. Preliminary results also indicated relatively high genomic prediction accuracy, suggesting that spatial correction, such as an AR1 × AR1 row column autocorrelation structure, would be unlikely to reduce error variance drastically. Instead, 59 plots that included breeder comments about bad seed or significant damage to the plot, via animal or otherwise, were removed from the dataset. Observations outside of a four standard deviation interval from the grand mean of uncorrected GY phenotypic observations were also removed to account for any significant undocumented damage, grain spillage or other undocumented mistakes. This included two observations that were deemed too high, and 20 observations that were deemed too low.

Observations of 11 lines lacking at least one phenotypic record in at least two separate trials and 60 lines that were missing at least 50% of their genotypic calls and one line that had greater than 20% heterozygous genotype calls were also removed from the dataset. This resulted in 8,692 phenotypic observations of 1,447 lines across 26 environments, representing 96.6% of the plots with grain yield measurements. HD was not recorded for the 246 observations from 2007, and PH was not recorded for the 840 observations from 2009. Two additional plots were missing PH measurements from the Ketola location, one that was recorded at 2 meters in 2008 which was likely a recording mistake, and one in 2010 which was simply missing a record.

**Figure A1:**
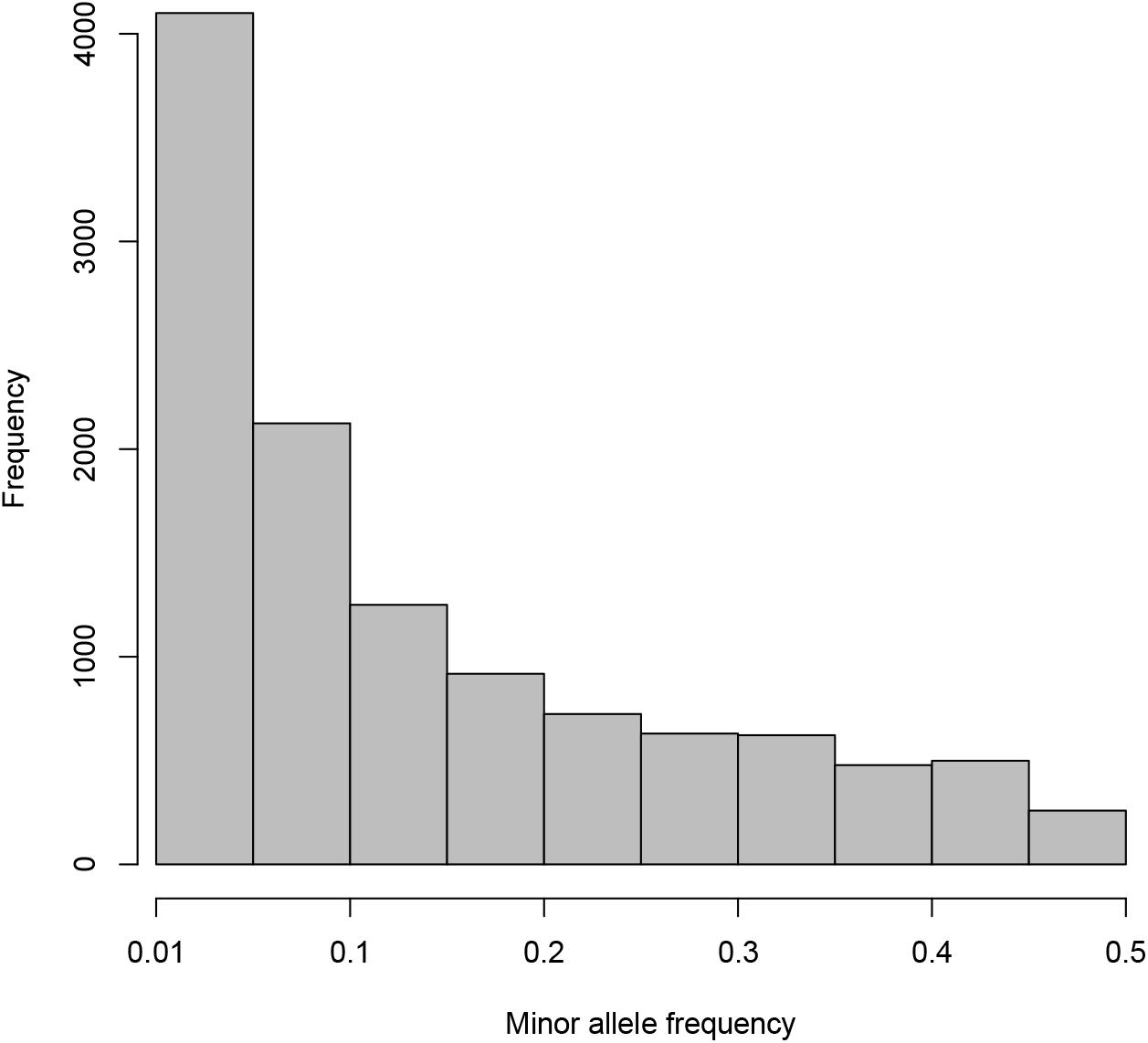
Distribution of minor allele frequencies for 11,604 GBS markers in the CNLM population.

While most of the genotypes were directly from the Cornell small grains breeding program, a few varieties and breeding lines from other breeding programs that had been genotyped and evaluated were not excluded from the dataset as long as they met the previous criteria. This included 75 lines from The Ohio State University wheat breeding program and 93 lines from the Michigan State University wheat breeding programs that were part of the Allele Based Breeding initiative, among other lines from various breeding programs.

Genotyping by sequencing (GBS) libraries (Elshire et al. 2011) of 1,521 CNLM were developed using the protocol described by Poland et al. (2012) at Kansas State University, and subsequently sequenced at the Genomic Diversity Facility at Cornell University. Genotyping calls were accomplished using standard parameters of the Tassel 5.0 GBS v2 Pipeline (Glaub-itz et al. 2014) and were aligned to the International Wheat Genome Sequencing Consortium (IWGSC) RefSeq v1.0 wheat genome sequence of ‘Chinese Spring’ (IWGSC 2018, accepted, unpublished). Following Poland et al. (Poland et al. 2012), 64 bp sequence tags containing no more than three Single Nucleotide Polymorphisms (SNPs) per tag were included to increase the likelihood of obtaining subgenome specific markers. Only markers with a minor allele frequency of at least 0.01 (Figure A1), no more than 30% missing scores, and no more than 10% heterozygous calls were kept for the following analyses. Then individuals with greater than 20% heterozygous calls and individuals with more than 50% missing genotype calls were excluded from the dataset. The process was repeated iteratively, starting by filtering on markers until the number of markers and genotypes converged. This resulted in 11,604 available GBS markers distributed throughout the three subgenomes (Figure A2). Of the 61 lines removed, 60 were removed due to missing marker information and 1 was removed due to high heterozygosity in the final iteration. The 11 lines with a single observation and the two lines without grain yield observations were subsequently removed to produce genotypic information for the 1,447 lines.

**Figure A2:**
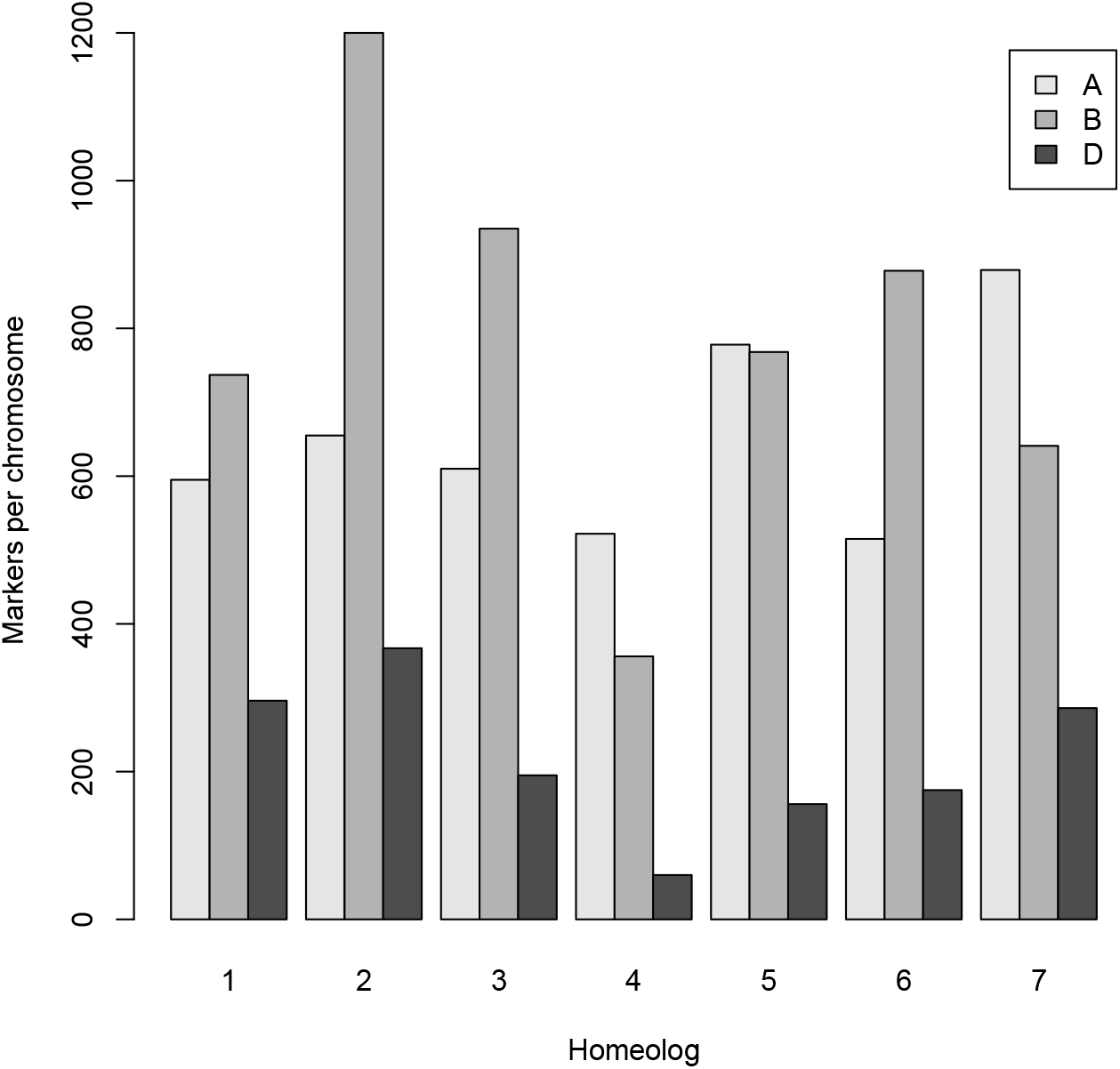
Distribution of 11,604 GBS markers on the 21 wheat chromosomes comprised of 7 homeologs of three subgenomes, A, B and D, for the CNLM population.

**Table A3:**
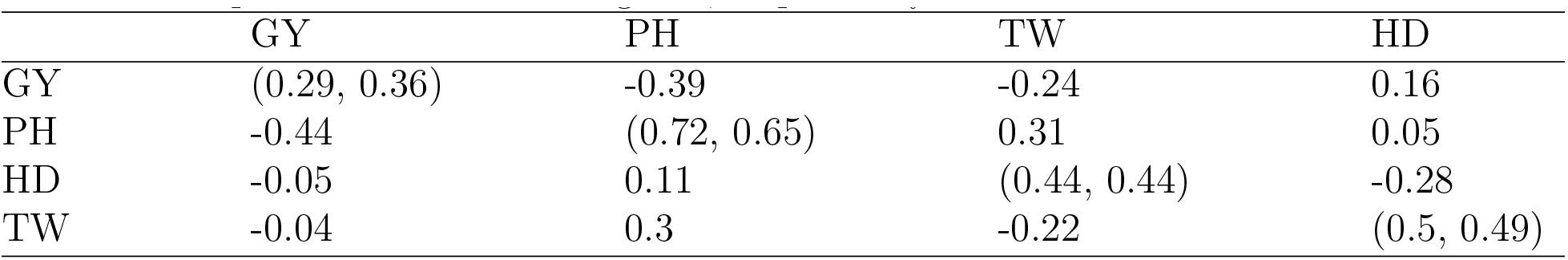
Estimated genetic correlation of traits with additive (below diagonal) and independent genetic relationships (above diagonal). Standard deviations of scaled traits estimated with a realized additive covariance between individuals and assuming independence are shown in parentheses on the diagonal, respectively.

Marker scores were coded using {−1,0,1} for homozygous major allele, heterozygous and homozygous minor allele, respectively. Categorical marker imputation was done independently for each chromosome using random forest imputation via the R package ‘missForest’ (Stekhoven and Buhlmann 2011) which relies on the R package ‘randomForest’ (Liaw and Wiener 2002). Random forest has been shown to be effective for genotype imputation in wheat (Rutkoski et al. 2013). To allow all individuals to be considered completely inbred, the remaining heterozygous calls (< 2% of all marker scores) were conservatively replaced with the population mode for that marker (i.e. the homozygous major allele, −1). Marker scores were then converted to {0,1} coding for presence of the minor allele.

Genetic correlations of traits were estimated in a multivariate model fit (A3). This was accomplished by treating genotypes as independent, or having a realized additive covariance structure calculated from genetic markers.

## A2 Appendix 2 Removal of population structure from M

Let **M** be the *n* × *m* matrix of *m* marker scores for *n* genotypes. Markers can be sorted into their respective genome, such that

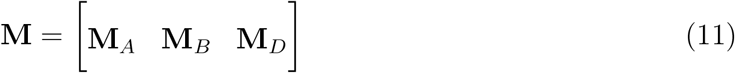

**M** can be factored using singular value decomposition as follows:

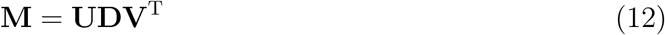

where **U**, and **V** are unitary matrices of left and right singular vectors, and **D** is a diagonal matrix of singular values.

The first *k* principal components of the marker matrix can be extracted by selecting the first *k* columns of **U** and the first *k* rows and columns of **D** and multiplying.

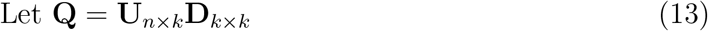

In a manner similar to Eckart and Young (Eckart and Young 1936), an approximation, 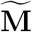, of the marker matrix, **M** with the first *q* principal components removed can be reconstructed by setting the first *k* singular values in **D** to zero (denoted 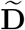).

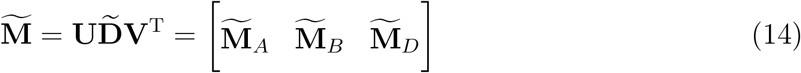

## S1 Supplementary Materials

**Figure S1:**
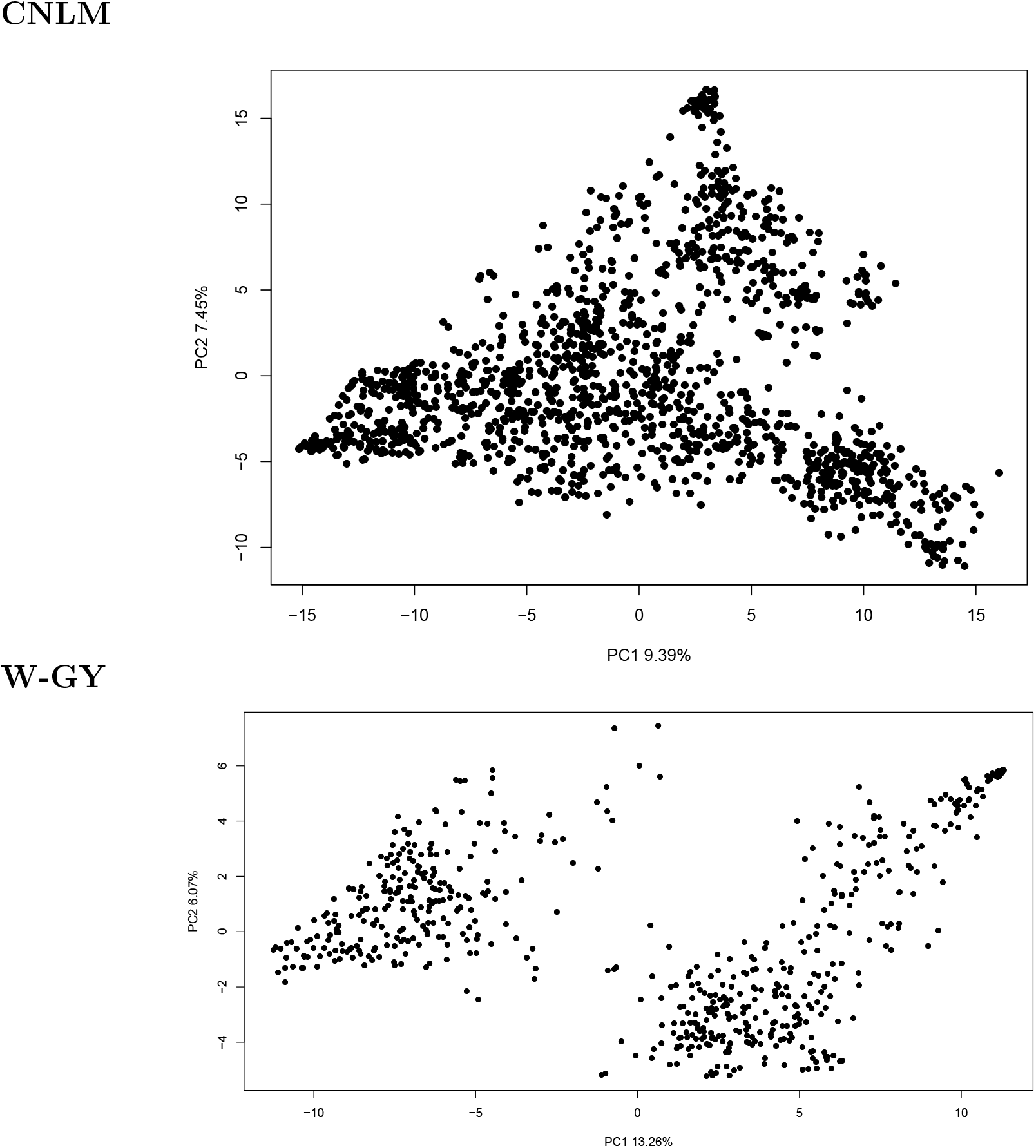
Plot of the first two principal components of the additive covariance matrix **K** in the CNLM and W-GY populations.

**Figure S2:**
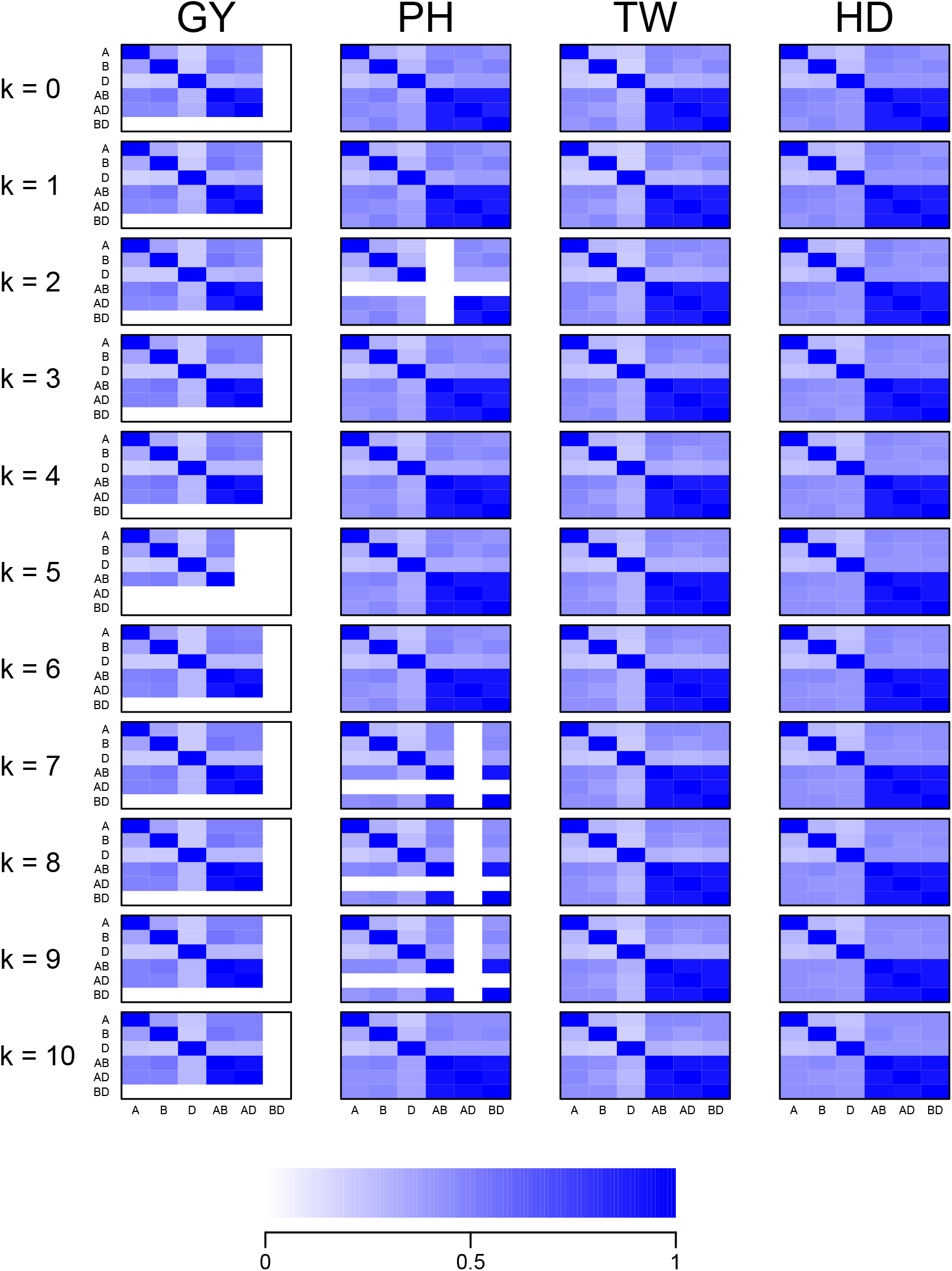
Correlation of variance component estimates derived from the average information from the model fit for models correcting for population structure with *k* ∈ {0,1,…, 10} principal components for four traits in the CNLM population.

**Figure S3:**
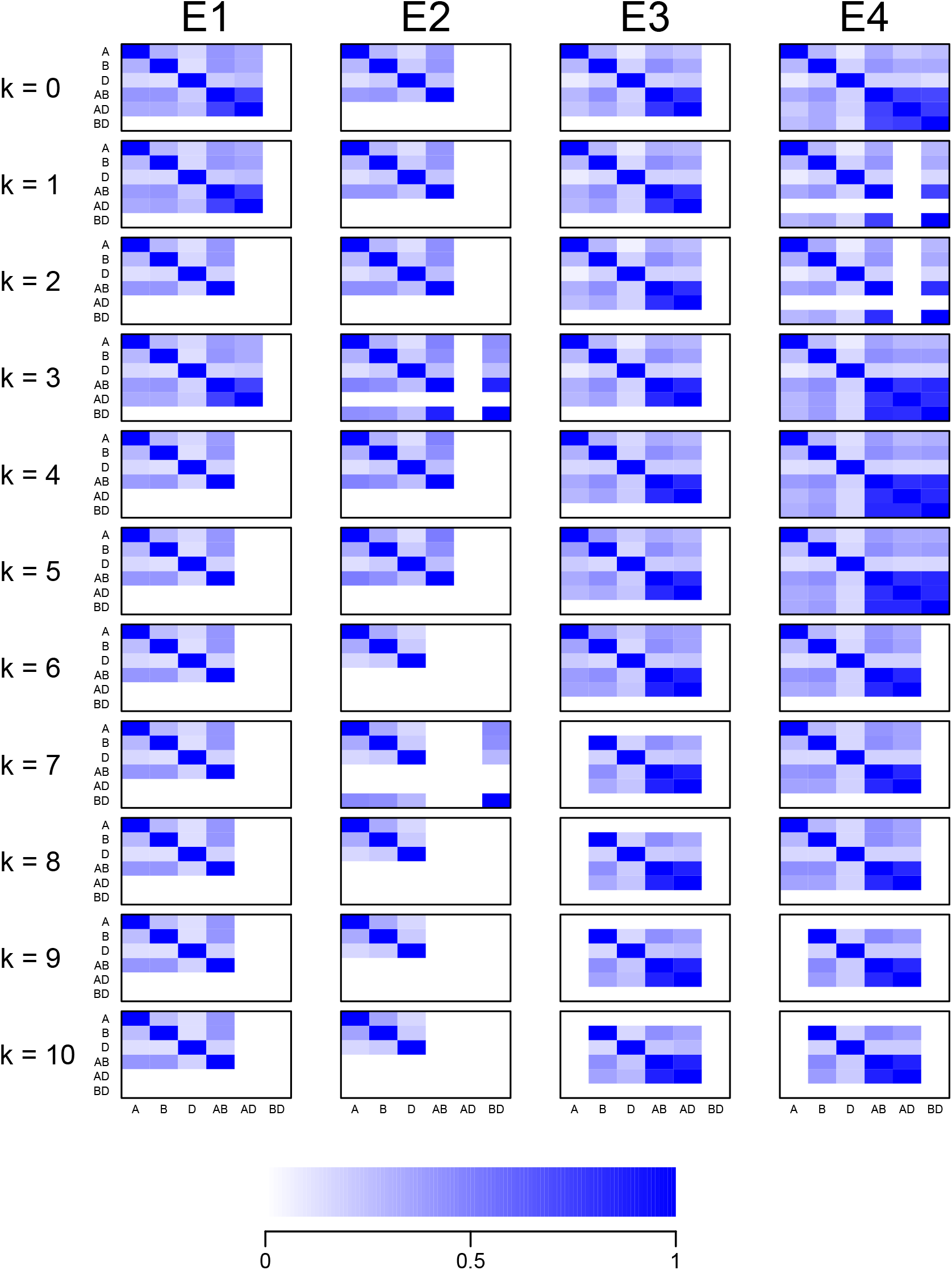
Correlation of variance component estimates derived from the average information from the model fit for models correcting for population structure with *k* ∈ {0,1,…, 10} principal components for four traits in the W-GY population.

## List of abbreviations

AIC: Akaike’s information criterion;
HD: heading date;
IWGSC: International Wheat Genome Sequencing Consortium;
GBS: genotyping by sequencing;
GEBV: genomic estimated breeding value;
GY: grain yield;
PC: principal component;
PH: plant height;
SGEBV: subgenomic estimated breeding value;
TW: test weight

